# Evolutionary rate coevolution between mitochondria and mitochondria-associated nuclear-encoded proteins in insects

**DOI:** 10.1101/288456

**Authors:** Zhichao Yan, Gongyin Ye, John H. Werren

**Author notes:** To whom correspondence should be addressed: John H. Werren, Department of Biology, University of Rochester, Rochester, NY 14627, USA., Gongyin Ye, Institute of Insect Sciences, C1152 Agro-Bio Complex, Zhejiang University, Hangzhou 310058, China.

## Abstract

The mitochondrion is a pivotal organelle for energy production, and includes components encoded by both the mitochondrial and nuclear genomes. How these two genomes coevolve is a long-standing question in evolutionary biology. Here we initially investigate the evolutionary rates of mitochondrial components (oxidative phosphorylation (OXPHOS) proteins and ribosomal RNAs) and nuclear-encoded proteins associated with mitochondria, across the major orders of holometabolous insects. There are significant evolutionary rate correlations (ERCs) between mitochondria and mitochondria-associated nuclear-encoded proteins, which is likely driven by different rates of mitochondrial sequence evolution and compensatory changes in the interacting nuclear-encoded proteins. The pattern holds after correction for phylogenetic relationships and considering protein conservation levels. Correlations are stronger for nuclear-encoded OXPHOS proteins in contact with mitochondrial-encoded OXPHOS proteins and nuclear-encoded mitochondrial ribosomal amino acids directly contacting the mitochondrial rRNA. Mitochondrial-associated proteins show apparent rate acceleration over evolutionary time, but we suspect this pattern to be due to artifacts (e.g. rate estimation or calibration bias). We find that ERC between mitochondrial and nuclear proteins is a strong predictor of nuclear proteins known to interact with mitochondria, and therefore ERCs can be used to predict new candidate nuclear proteins with mitochondrial function. Using this approach, we detect proteins with high ERCs but not with known mitochondrial function based on gene ontology (GO). Manual screening of the literature revealed potential mitochondrial function for some of these proteins in humans or yeast. Their holometabolous ERCs therefore indicate these proteins may have phylogenetically conserved mitochondrial function. Twenty three additional candidates warrant further study for mitochondrial function based on this approach, including ERC evidence that proteins in the minichromosome maintenance helicase (MCM) complex interact with mitochondria. We conclude that the ERC method shows promise for identifying new candidate proteins with mitochondrial function.

## Introduction

The mitochondrion is a product of an ancient symbiosis (Pittis & Gabaldon 2016; Sagan 1967), and despite considerable stream-lining and gene loss relative to its bacterial ancestor, still contains a subset of its own genes, especially those in the oxidative phosphorylation pathway (OXPHOS or OP), ribosomal RNA, and some tRNAs (Rand 2008; Rand *et al.*, 2004). Many other proteins are encoded in the nucleus and transported to the mitochondria. These include a large number of nuclear-encoded mitochondrial ribosomal proteins (**nMRPs**) and protein components of the OXPHOS pathway (**nOPs**). As a result, there are extensive molecular interactions between the products of mitochondria and mitochondrial associated nuclear-encoded proteins (Gray 2012; Hill 2015; Rand *et al.*, 2004). Mitochondrial-nuclear incompatibilities can be lethal or severely reduce fitness in species hybrids in some taxa, suggesting its potential contribution to the process of speciation (Arnqvist *et al.*, 2010; Bolnick *et al.*, 2008; Breeuwer & Werren 1995; Burton *et al.*, 2013; Chou & Leu 2010; Ma *et al.*, 2016; Meiklejohn *et al.*, 2013; Niehuis *et al.*, 2008; Rawson & Burton 2002). For example, an important role of nuclear-cytoplasm incompatibility in hybrids fitness reduction has been found in several systems, including *Nasonia* wasps (Breeuwer & Werren 1995; Niehuis *et al.*, 2008), whereas cyto-nuclear incompatibility appears to be less important to hybrid breakdown in other taxa, such as *Drosophila* (Meiklejohn *et al.*, 2013; Montooth *et al.*, 2010). The direct interactions between gene products from different genomic compartments raises fascinating questions about how nuclear and mitochondrial genomes coevolve, especially in light of different substitution rates among nuclear and mitochondrial genomes in taxa (Hill 2015; Rand *et al.*, 2004).

Although mitochondria share a single origin from endosymbiotic integration of an α-Proteobacteria (Pittis & Gabaldon 2016; Sagan 1967), eukaryotes have quite divergent substitution rates in mitochondrial genomes, even in some closely related species (*Baer et al.*, 2007; Lynch *et al.*, 2006; Sloan *et al.*, 2009). An extreme difference is between animals and plants. Mitochondrial substitution rates in animals are ~100 times higher than plants (*Lynch et al., 2006*; Palmer & Herbon 1988), and there is considerable variation in rates within plants and animals (Grossman *et al.*, 2004; Oliveira *et al.*, 2008; Sloan *et al.*, 2012; Sloan *et al.*, 2009). In contrast, substitution rates in nuclear genomes are similar between animals and plants (Lynch 2010). The higher substitution rates in animal mitochondria are believed to be driven by elevated mutation rates compared to nuclear genomes. In insects, mitochondrial genomes from Hymenoptera and Psocodea show exceptionally high rates of substitution and gene order rearrangement whereas rates are much lower in Diptera (Cameron 2014; *Kaltenpoth et al.*, 2012b; Oliveira *et al.*, 2008; Yoshizawa & Johnson 2013). For example, the synonymous substitution rate for mitochondrial DNA in the parasitoid wasp *Nasonia* is greater than 40 times higher than the nuclear genes (Oliveira *et al.*, 2008). However, the causes and consequences of this rate variation are still largely unknown (Baer *et al.*, 2007).

It has also been observed that rates of mitochondrial evolution appear to be accelerating across evolutionary time (Ho *et al.*, 2005; Molak & Ho 2015). This apparent acceleration has been observed in metazoan mitochondria across a wide range of time and taxa. However, the pattern may be due to calibration errors or rate estimation bias, mainly due to saturation in long branches and standing variation at the “tips” of terminal branches (e.g. caused by incomplete purifying selection), creating the appearance of lower rates in the past (Ho *et al.*, 2011; Ho *et al.*, 2015).

In contrast to plants, animals contain streamlined mitochondrial genomes with substitution rates commonly higher than nuclear genomes (Barreto & Burton 2013; Grossman *et al.*, 2004; Lin & Danforth 2004; Mishmar *et al.*, 2006; Werren *et al.*, 2010; Willett & Burton 2004). Several theories are proposed to explain this, such as cell cycle independent replication in mitochondria, increased mutagenic pressure of oxidative radical species, and limited DNA repair ability (Neiman & Taylor 2009). As mitochondria are maternally inherited without recombination, the effective population size is relatively small and it is more likely to accumulate mildly deleterious mutations by genetic drift. Hitchhiking of mildly deleterious mutations is also more likely to occur in the non-recombining genome of mitochondria. Furthermore, because of maternal inheritance, the mitochondrial genome is under no selection in males. Mutations only deleterious to males could be maintained and reach a high frequency, which is known as “mother’s curse” (Gemmell *et al.*, 2004; Hill 2015; Innocenti *et al.*, 2011). To alleviate the effect of mitochondrial deleterious mutations, it is also proposed that adaptive compensatory nuclear mutations are likely, and nuclear adaptive selection will be accelerated by recombination and relatively large population size (Barreto & Burton 2013; Breeuwer & Werren 1995; Oliveira *et al.*, 2008; Osada & Akashi 2012; Sloan *et al.*, 2014). In addition, a feedback can occur when compensatory mutations produce pleiotropic effects that select for additional adaptive mitochondrial substitutions (Rand *et al.*, 2004). Such events can lead to further fixation of mildly deleterious mitochondrial mutations by hitchhiking during the adaptive sweep, setting up a cycle of nuclear compensation – mitochondrial compensation and deleterious draft – nuclear compensation (Oliveira *et al.*, 2008). The rate of feedback would be accelerated in taxa with elevated mitochondrial mutation rates. For these various reasons, nuclear-mitochondrial interactions are expected to promote coevolution of mitochondrial and nuclear genomes. Nevertheless, few studies have investigated coevolution of rates of mitochondrial and nuclear encoded proteins, except in a plant lineage *Silene* (Havird *et al.*, 2017; Havird *et al.*, 2015; Sloan *et al.*, 2014), where mitochondrial evolving rates are generally lower than those in animals, and a recent study of evolutionary rate correlations in OXPHOS genes of insects, with an emphasis on Hymenoptera (Li *et al.*, 2017).

Here we investigate the evolutionary pattern of mitochondria-associated nuclear-encoded proteins in holometabolous insects, and their rate coevolution with mitochondrial-encoded proteins and ribosomal RNAs. In summary, we show 1) positive correlations in evolutionary rates of mitochondria and mitochondria-associated nuclear proteins; 2) even stronger rate correlations for OXPHOS proteins that make physical contact with mitochondrial OXPHOS proteins, and amino acids from nuclear-encoded MRPs contacting the mitochondrial rRNAs or tRNAs; 3) that nOP and nMRP have different patterns from other nuclear proteins over evolutionary time; 4) this creates the appearance of their accelerating over evolutionary time, but that this is likely an artifact; and finally 5) that evolutionary rate correlation (ERC) is a powerful predictive tool for mitochondria interacting nuclear proteins, and can be used to predict new nuclear protein candidates with mitochondrial function.

## Results & Discussion

### 1. Rate coevolution of mitochondrial products and mitochondria-associated nuclear proteins

To investigate the coevolution between mitochondrial products and nuclear-encoded proteins, we initially use protein data and the phylogenetic tree topology published in the paper on insect phylogeny from Misof *et al.* (2014), which was constructed using 1478 single copy nuclear genes. For each 64 holometabolous taxa in that analysis, we identified available mitochondrial genomes from either the same insect species or a relative (see methods, Table S1). We did not attempt to extract mitochondrial genomes from insect transcriptomic data, as has been used in some studies (Li *et al.*, 2017; Song *et al.*, 2016), because our initial evaluations suggested contamination problems, possibly from nuclear sequences of mitochondrial origins and/or barcode index hopping (Ballenghien *et al.*, 2017; Illumina 2017).

Evolutionary rate correlation (**ERC**, abbreviations see Table 1) between mitochondria-associated nuclear proteins and mitochondrial components were investigated in holometabolous insects. We defined the following categories for mitochondrial-encoded products and nuclear-encoded proteins. For mitochondrial-encoded components, 13 mitochondrial encoded oxidative phosphorylation proteins (**mOP**) and mitochondrial 12S and 16S ribosomal RNA (**mrRNA**) were utilized. From the 1478 single copy nuclear-encoded proteins retrieved from Misof *et al.* (2014), 19 nuclear-encoded OP proteins (**nOP**), and 26 nuclear-encoded mitochondrial ribosomal proteins (**nMRP**) were utilized. Using the Gene Ontology (Falcon & Gentleman 2007; Gramates *et al.*, 2017) and MitoDrome (D’Elia *et al.*, 2006) data bases (see Methods), we identified a set of 138 mitochondria-associated proteins, which includes our nOP and nMRP proteins. These 138 mitochondria-associated proteins were removed from the 1478 single copy protein set to generate a control set of 1340 proteins (**nALL\***) for comparative analyses. An additional nuclear encoded protein category is defined that serves as a control, 29 cytosolic ribosomal proteins (**nCRP**). The nCRPs provide a useful comparison in nMRPs because both sets of proteins are associated with ribosomal RNA, in one case the nuclear encoded ribosomes, and in the other case the mitochondrial encoded ribosome. Initially, we calculated branch lengths for the concatenated proteins in each gene set, which represent numbers of substitutions per site based on the tree topology (branch pattern) for the relevant taxa (Misof *et al.*, 2014). Later analyses (see below) calculated substitution patterns for individual proteins.

**Table 1.**
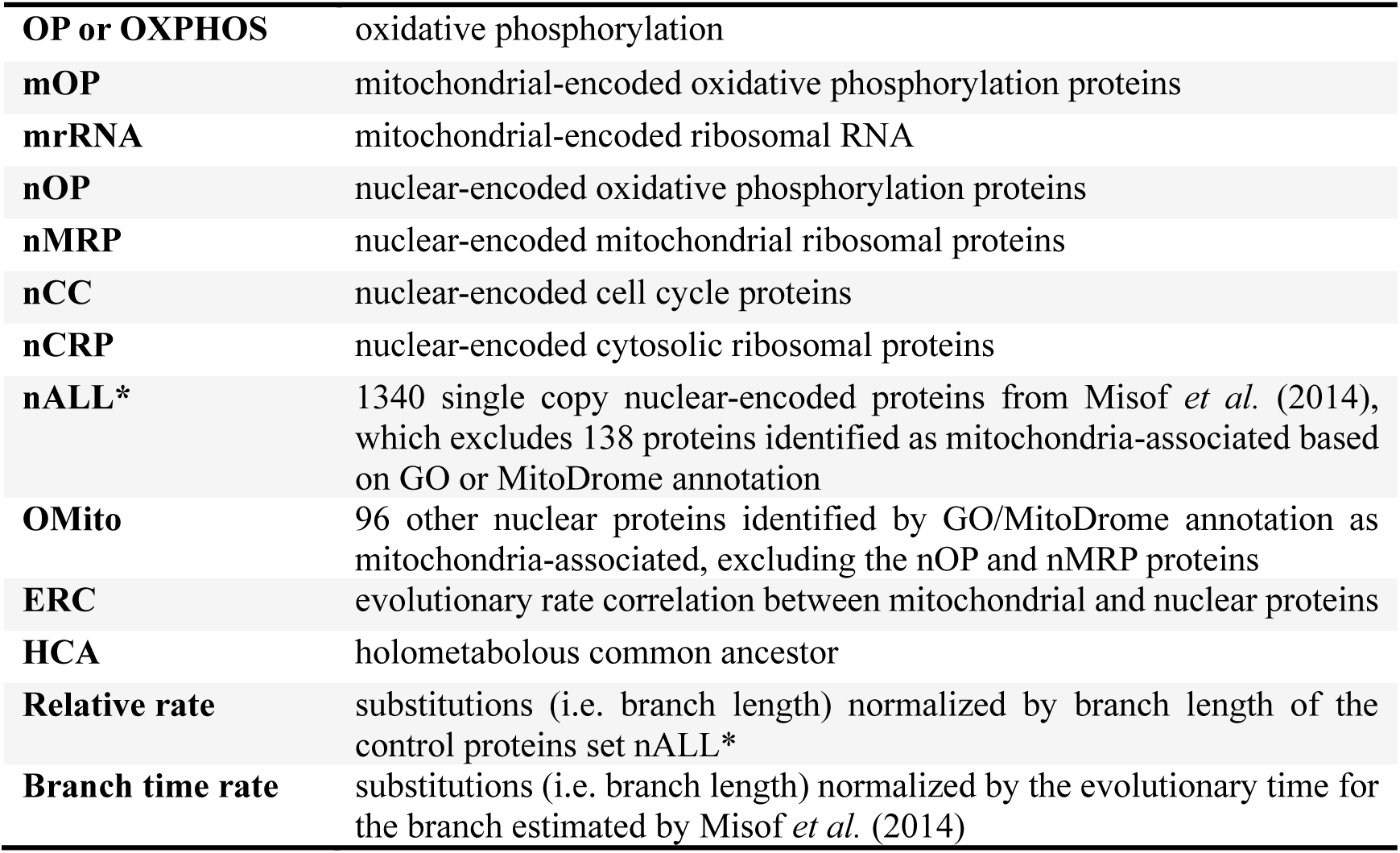
Abbreviations used in this paper.

Figure 1 shows the branch lengths from the holometabolous common ancestor (**HCA**) for concatenated sets of mitochondrial RNAs (mRNA), mitochondrial OPs (mOP) and nuclear protein categories (nOP, nMRP, nCRP, and nALL*) in 11 holometabolous insect orders. As can be seen, there are considerable differences in protein branch lengths between the different orders, and between the gene categories. Statistical groupings of the orders determined by Fisher’s least significant difference followed by Benjamini & Hochberg multiple correction (Benjamini & Hochberg 1995) are also shown. As already reported in other studies (Kaltenpoth *et al.*, 2012a; Li *et al.*, 2015; Li *et al.*, 2017), the Hymenoptera have significantly longer branches (faster rates of evolution) than most other orders in mOP and mrRNA, with Strepsiptera also showing elevated rates. In contrast, the Diptera show relatively slower rates of evolution in mitochondrial OPs and rRNAs (details in supplementary materials section S1).

**FIGURE 1.**
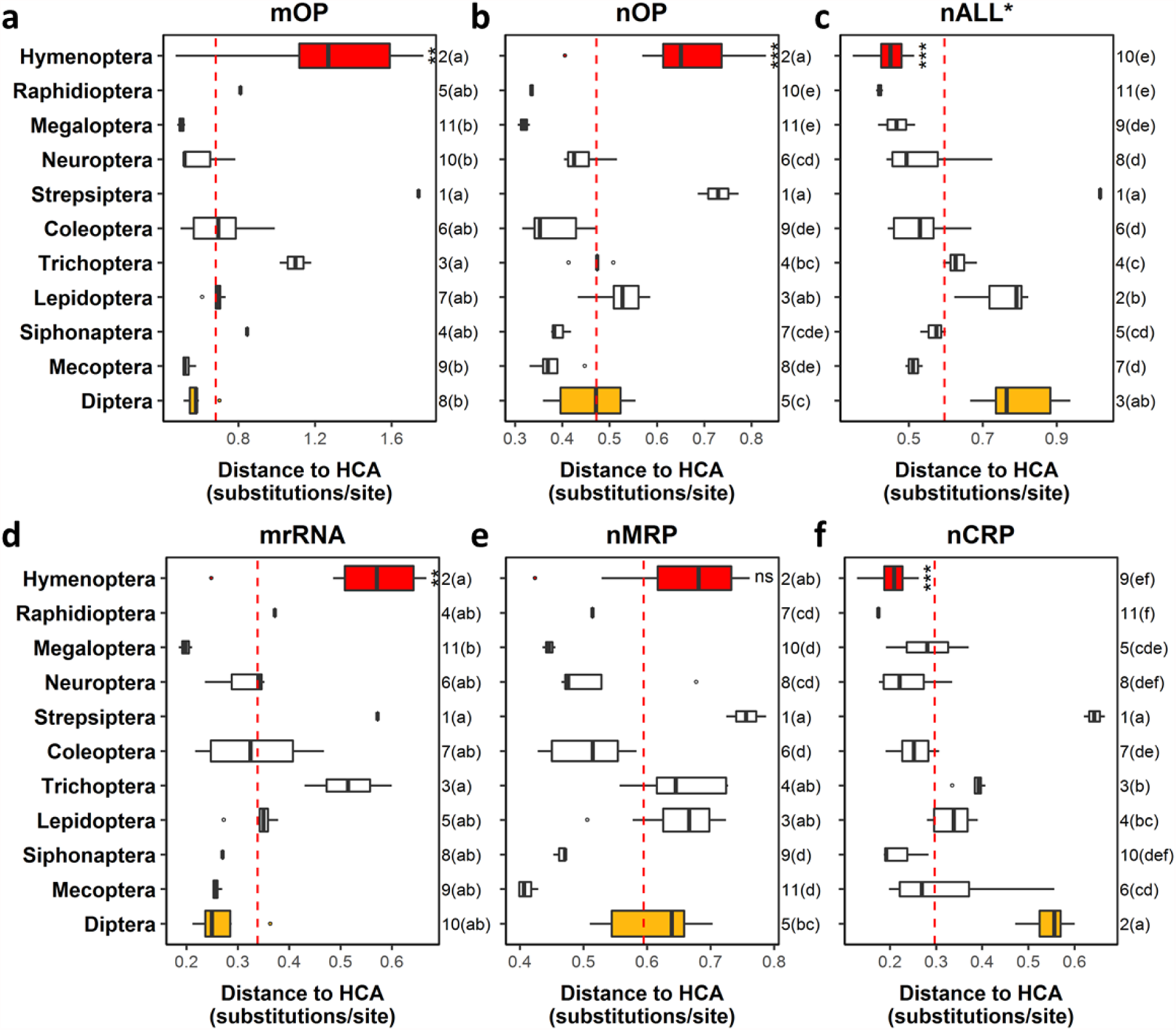
Distances to the holometabolous common ancestor (HCA) of concatenated mitochondrial products and nuclear protein categories for different orders. Distance to HCA are shown for **(a)** mitochondrial-encoded OXPHOS proteins (**mOP**), **(b)** nuclear-encoded OXPHOS proteins (**nOP**), **(c)** all 1478 nuclear-encoded proteins excluding these with mitochondrial GO annotations (**nALL\***), **(d)** mitochondrial ribosomal RNAs (**mrRNA**), **(e)** nuclear-encoded mitochondrial ribosomal proteins (**nMRP**), and **(f)** nuclear-encoded cytosolic ribosomal proteins (**nCRP**). For mitochondrial products, 44 species with associated mitochondrial genomes were used (see methods). For nuclear-encoded proteins, 64 holometabolous species were included for stronger statistical power and a more comprehensive analysis. Numbers on right indicate ranks of each order. Orders sharing the same letter(s) in parentheses have no significant difference from each other (*p* < 0.05). Asterisks on Hymenoptera indicate significant difference from Diptera using Wilcoxon rank sum test; ns indicates no significant difference (*p* > 0.05); * *p* < 0.05; ** *p* < 0.01; *** *p* < 0.001.

To compare rates of evolution in mitochondrial and nuclear components, we first conducted an evolutionary rate correlation (ERC) using a non-parametric Spearman (NPS) approach between mitochondrial and nuclear distances to the holometabolous common ancestor (**HCA**). Distances to HCA of mOP show significant correlation with those of nOP (Figure S1; Spearman’s *ρ* = 0.70, *p* = 3.5e^−7^). Similarly, distances to HCA of mrRNA show positive correlation with those of nMRP (Figure S1; *ρ* = 0.66, *p* = 2.1e^−6^). In contrast, there is no significant correlation between mOP and the control protein set nALL* (*p* = 0.99). Nor do the control proteins nALL* (*p* = 0.72) and nCRP (*p* = 0.38) show significant rate correlations with mrRNA.

The patterns above are consistent with evolutionary rate correlations between mitochondrial components and interacting nuclear components such as nOP and nMRP. However, as distances to HCA share internal branches, they are not independent measures of evolutionary rates for each taxon within the tree. We therefore next investigated nuclear-mitochondrial correlations using evolutionary rates on terminal branches (Figure 2). Two different methods were used for calculating rates, either normalization to branch length of the control protein set nALL* (henceforth referred to as the **relative rate**) or normalization to the evolutionary time of the terminal branch estimated by Misof *et al.* (2014) (henceforth referred to as the **branch time rate**).

**FIGURE 2.**
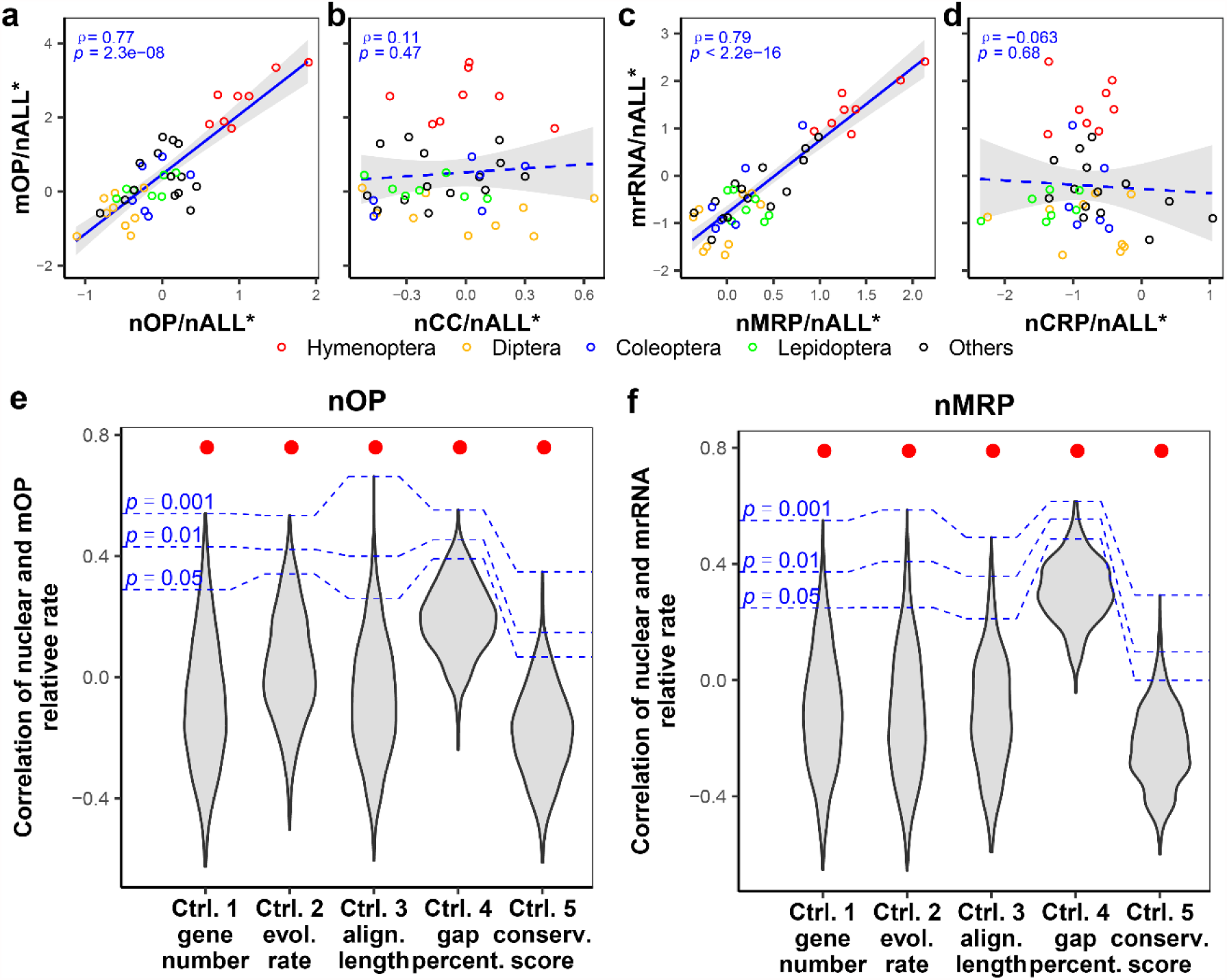
Relative rates coevolve between mitochondrial products and mitochondria-associated nuclear proteins. **(a–d)** Correlations between relative rates of **(a)** mitochondrial and nuclear-encoded OXPHOS proteins (**mOP** and **nOP**), **(b)** mOP and nuclear-encoded cell cycle proteins (**nCC**), **(c)** mitochondrial rRNA (**mrRNA**) and mitochondrial ribosomal proteins (**nMRP**), **(d)** mrRNA and nuclear-encoded cytosolic ribosomal proteins (**nCRP**). Relative rates in **a–d** were log2 transformed for data visualization. Spearman’s ρ and *p* value are presented. A solid regression line indicates a significant Spearman correlation, and dashed regression line a non-significant correlation. Gray shadows represent 95% confidence intervals for regression lines. **(e–f)** Subsampling statistics are shown for **(e)** nOP-mOP relative rate evolutionary correlation (**ERC**) and **(f)** nMRP-mrRNA ERC. Subsampling was conducted by randomly sampling from nALL* proteins. Five different methods were used to control for either gene number, evolutionary rate, alignment length, gap percentage, or conservation score. Red dots indicate the observed ERC of nOP-mOP or nMRP-mrRNA. As can be seen, ERC of nOP-mOP and nMRP-mrRNA are significantly higher than ERC of mitochondrial products and randomly generated samples.

Relative rate estimates are shown in Figure 2a–d. The relative rate estimated for nOP are strongly positively correlated with mOP (*ρ* = 0.77, *p* = 2.3e^−8^), and relative rate of nMRP also strongly positively correlate with mrRNA (*ρ* = 0.79, *p* < 2.2e^−16^). However, nCRP and mrRNA are not significantly correlated (*p* = 0.68). An alternative control, nuclear-encoded cell cycle proteins (**nCC**) (methods), is used and shows no significant correlation with mOP (*p* = 0.47). Relative rates for nALL* are not shown because nALL* is used for rate normalization. Branch time normalization allows for rate estimates of nALL*. Branch time rates of mOP show strongly significant correlation with nOP (Figure S2; *ρ* = 0.71, *p* = 2.3e^−7^), but no correlation with nALL* (*p* = 0.16) or nCC (*p* = 0.14); mrRNA show strong correlation with nMRP (*ρ* = 0.78, *p* = 9.8e^−9^), but no correlations with nALL* (*p* = 0.14) or nCRP (*p* = 0.52).

The results above are consistent with nOPs and nMRPs having significant evolutionary rate correlations with corresponding mitochondrial components. However, it could be argued that the patterns are dominated by certain taxa (e.g. the Hymenoptera) and do not broadly occur across the holometabolous insects. When we perform a regression analysis using insect order as a variable, the same basic patterns were confirmed (statistical details see Table S2). Similarly, if we remove Hymenoptera from the analysis, there are still significant positive evolutionary rate correlations between nOP and mOP (Table S2; relative rate: *ρ* = 0.58, *p* = 0.0003; branch time rate: *ρ* = 0.60, *p* = 0.00014) and between nMRP and mrRNA (relative rate: *ρ* = 0.63, *p* = 6.5e^−5^; branch time rate: *ρ* = 0.71, *p* = 2.5e^−6^). Finally, we also applied a non-parametric independent contrasts test (Garland *et al.*, 1992). Results confirm strong significant relative rate correlations between nOP-mOP (Figure S3; *ρ* = 0.53, *p* = 0.00021) and nMRP-mrRNA (Figure S3; *ρ* = 0.77, *p* = 1.4e^−9^), and similar results hold for branch time evolutionary rate (Figure S4).

To further rule out potential artifacts, subsampling analyses (Politis *et al.*, 1999) were conducted for differences between concatenate mitochondria-associated and control protein sets, including gene number, alignment length, evolutionary rate, conservation score and gap percentage. Basically the method subsamples from the nALL* control protein set to produce 1000 independent control samples for each feature. The ERC distribution of each control set is then used to generate the null hypothesis statistical expectation for the experimental set (e.g. nOP, nMRP). Figure2e–f compares the control ERC distributions using relative rates on terminal branches to the observed mitochondria-associated proteins for the different features. For these features, both nOP and nMRP show significantly higher relative rate and branch time rate ERCs than randomly generated controls (Figure 2e–f, Figure S5; *p* < 0.001 in each case). Therefore, the ERCs between mitochondria-associated nuclear proteins and mitochondria are unlikely to be due to unusual features of the protein sets. Furthermore, although we didn’t control amino acid composition of nOPs and nMRPs by subsampling, because of its high dimensions with 20 amino acids, comparisons of amino acid composition indicates that they are not dramatically different to control proteins for nOP and nMRP (Figure S6).

To have a more comprehensive understanding of ERC, individual gene analyses were also conducted. After model testing for the most appropriate evolutionary model per protein (Stamatakis 2014), relative rates for terminal branches of each protein were computed (Table S3). Results (Figure 3, Table S4) show that nOP have significantly higher ERC to mOP than do nALL* and nCC (cell cycle proteins), for both relative and branch time rates (Wilcoxon rank sum test (WRST): *p* < 0.001), as do nMRP with mrRNA compared to nCRP and nALL* (WRST: *p* < 0.001). These results are consistent with the previous results using concatenated alignments. We also observed that both relative and branch time rates of **OMito** (other nuclear proteins with mitochondrial gene ontology) show significantly higher ERC with mitochondrial products than do nALL* (Table S4; WRST: *p* < 0.001).

**FIGURE 3.**
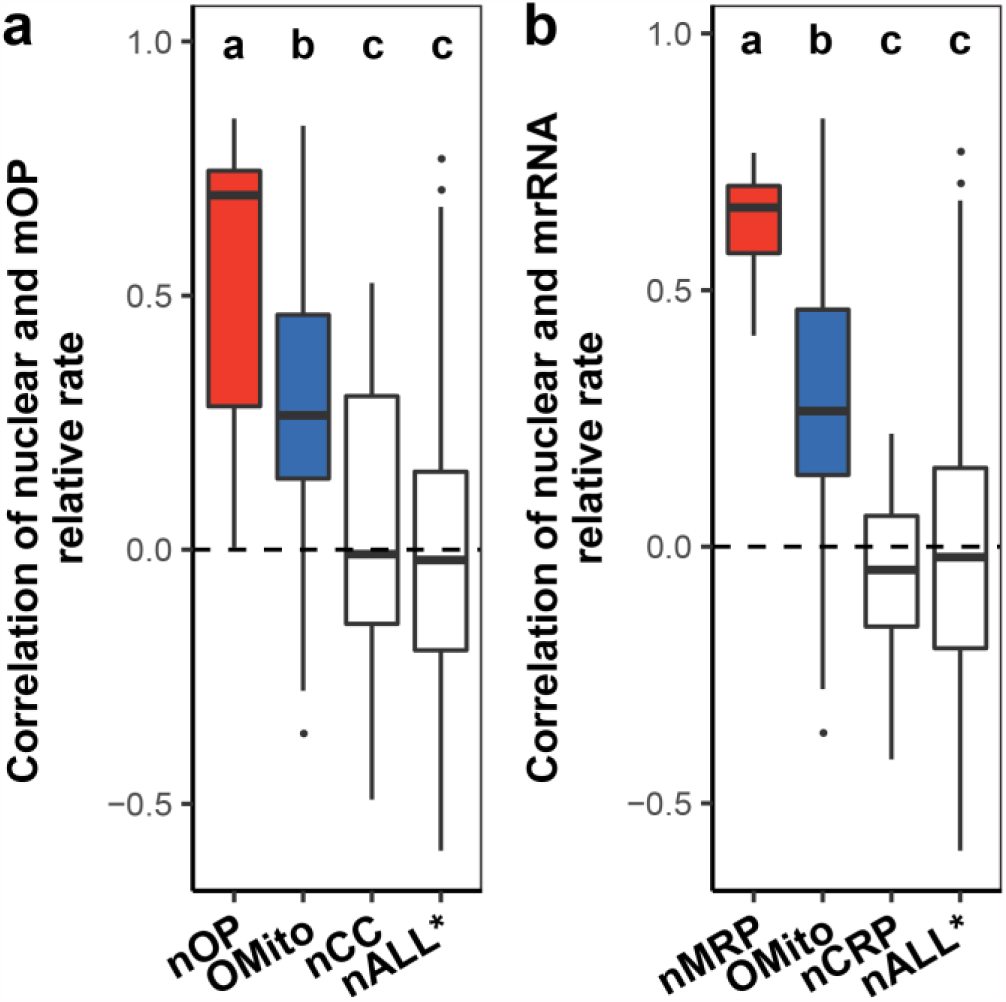
Comparison of evolutionary rate correlations between mitochondrial products and different nuclear gene categories using individual gene analyses. **(a)** Shown are correlations of relative rates of mOP to **nOP** (nuclear-encoded oxidative phosphorylation proteins), **OMito** (other nuclear-encoded proteins with mitochondrial annotations excluding nOP and nMRP), **nCC** (nuclear-encoded cell cycle proteins) and **nALL\*** (without mitochondrial gene annotation). Also shown are **(b)** correlations of relative rates of mrRNA to **nMRP** (nuclear-encoded mitochondrial ribosomal proteins), OMito, **nCRP** (nuclear-encoded cytosolic ribosomal proteins), and nALL*. Significant differences were determined by pairwise Wilcoxon rank sum test among protein categories, and followed by Benjamini & Hochberg multiple corrections. Groups sharing the same letter(s) have no significant difference from each other (*p* < 0.05). As seen, mitochondria-associated proteins show higher relative evolutionary rate correlations with mitochondrial products than do other nuclear-encoded proteins (*p* < 0.01 for each case).

An independent case of highly elevated rates of mitochondrial evolution and associated rapid evolution of mitochondrial-associated nuclear proteins was also found within Psocodea (supplementary section S2). In summary, the results indicate strong evolutionary rate correlations between mitochondria and mitochondria-associated nuclear encoded proteins.

### 2. Directly interacting nuclear proteins and amino acids in contact with mitochondrial components show stronger nuclear-mitochondrial coevolution

We investigated whether physical contacts with mitochondrial components contribute to evolution of mitochondria-associated nuclear-encoded proteins. Knowledge of the structures of mammalian mitochondrial complexes allow us to separate the mitochondria-targeted nuclear genes into components directly in contact with mitochondrial encoded components and those not. Based on structure information, nuclear coded OP proteins were divided into those directly in contact with mitochondrial encoded proteins and those not. Results confirm that nOPs in direct contact with mOPs have higher ERCs with mOP than do those not in contact. Terminal evolutionary rates of nOPs in contact with mOP are positively correlated with those of mOP (Figure 4; relative rate: *ρ* = 0.79, *p* < 2.2e^−16^; Figure S7; branch time rate: *ρ* = 0.83, *p* < 2.2e^−16^), but nOPs not in contact with mOP are not (relative rate: *p* = 0.12; branch time rate: *p* = 0.35). As almost all nMRPs directly contact mrRNA, we separated their individual amino acids into those in contact with mitochondrial rRNA or tRNA and those not (Figure S8). Both evolutionary rate of nMRP amino acids in contact (Figure 4; relative rate: *ρ* = 0.79, *p* = 5.3e^−10^; Figure S7; branch time rate: *ρ* = 0.82, *p* < 2.2e^−16^) and those not in contact (relative rate: *ρ* = 0.74, *p* = 9.5e^−8^; branch time rate: *ρ* = 0.71, *p* < 2.4e^−7^) with mitochondrial rRNA/tRNA show positive correlation with mrRNA. nMRP amino acids in contact with mitochondrial rRNA/tRNA show slightly but significantly higher ERC with mrRNA than do those not contact (dependent Spearman correlations comparison (DSCC) (Rosner *et al.*, 2015); relative rate: *p* = 0.046; branch time rate: *p* = 0.008).

**FIGURE 4.**
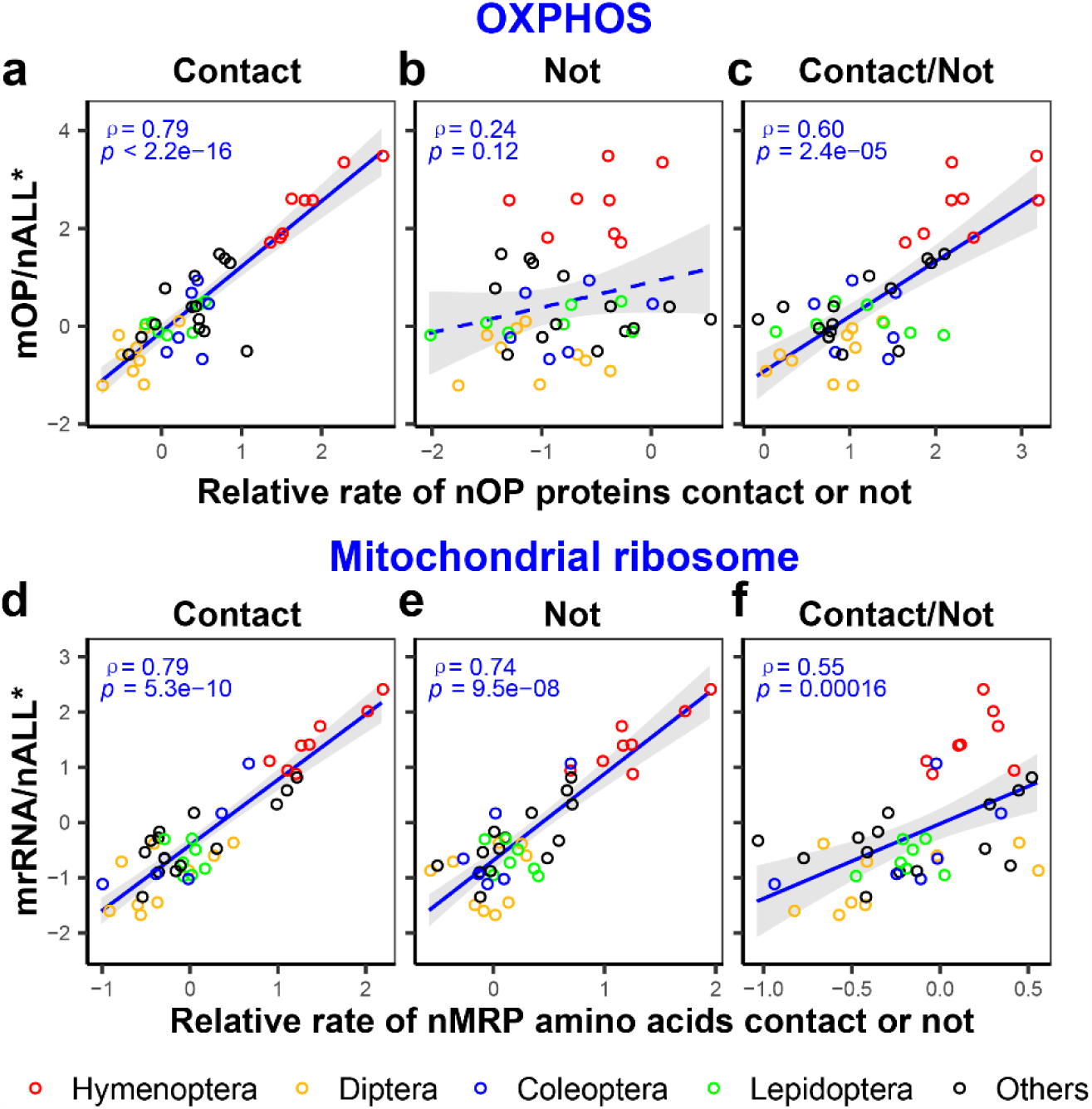
Mitochondria-associated nuclear proteins in contact with mitochondrial products show stronger mitochondrial-nuclear coevolution. Shown are correlations between relative rates of mOP and **(a)** nOP proteins in contact with mOP, **(b)** nOP proteins not in contact with mOP and **(c)** evolutionary rate ratio of contact over not contact of nOP. Also shown are correlation between relative rates of mrRNA and **(d)** nMRP amino acids in contact with mitochondrial rRNA/tRNA, **(e)** nMRP amino acids not in contact with mitochondrial rRNA/tRNA and **(f)** evolutionary rate ratio of contact over not contact amino acids for nMRP. Relative rates and ratios were log2 transformed. Spearman’s *ρ* and p value are presented. A solid regression line indicates a significant Spearman correlation (*p* < 0.05), and dashed regression lines a non-significant correlation. Gray shadows represent 95% confidence intervals for regression lines.

We further calculated evolutionary rate ratios of contacting proteins (nOP) to those not in contact. Similarly, contacting nMRP amino acids were compared relative to non-contacting nMRP amino acids. The contact over not contact rate ratio of nOP shows a significant positive correlation with both relative and branch time rates of mOP (Figure 4; relative rate: *ρ* = 0.60, Figure S7; *p* = 2.4e^−5^; branch time rate: *ρ* = 0.59, *p* = 3.1e^−5^). The contact over not contact ratio of nMRP amino acids also shows a positive correlation with both relative and branch time rates of mrRNA (relative rate: *ρ* = 0.55, *p* = 0.00016; branch time rate: *ρ* = 0.65, *p* = 3.3e^−6^). Correlations in this section remain significant after independent phylogenetic contrasts (Figure S9–10), except for a nearly significant correlation between rate ratio of contact over not contact nMRP amino acids and mrRNA (*p* = 0.079). In addition, individual protein analyses also show that nOP in contact with mOP show higher correlations than do those not in contact for both rate methods (Table S4; WRST; *p* = 0.002).

These results indicate that nuclear-encoded proteins or amino acid sites that contact with mitochondrial-encoded products have stronger ERC (coevolution) with mitochondrial-encoded products than those that are not in contact.

### 3. Apparent acceleration of nuclear proteins associated with mitochondria

Metazoan mitochondria appear to show acceleration in their rate of change over evolutionary time (Ho *et al.*, 2005; Molak & Ho 2015). However, the pattern is likely an artifact generated by incomplete purifying selection on terminal branches and substitution saturation in deep branches (Molak & Ho 2015). We conducted an extensive analysis of time-dependent evolutionary pattern of mitochondria-associated nuclear proteins (nOPs and nMRPs) relative to those without mitochondrial annotations. Details are presented in supplemental materials (section S3) and summarized here. We focus our attention on branch time rates because this allows a comparison of evolutionary rate patterns for mitochondria-associated versus other nuclear proteins. There are striking differences between mitochondria-associated proteins (nOP and nMRP) and other nuclear proteins in the patterns of their rates of change over evolutionary time (Figure 5). Subsampling results indicate that saturation cannot easily explain apparent rate acceleration for the mitochondria-associated nuclear proteins (Figure S11).

**FIGURE 5.**
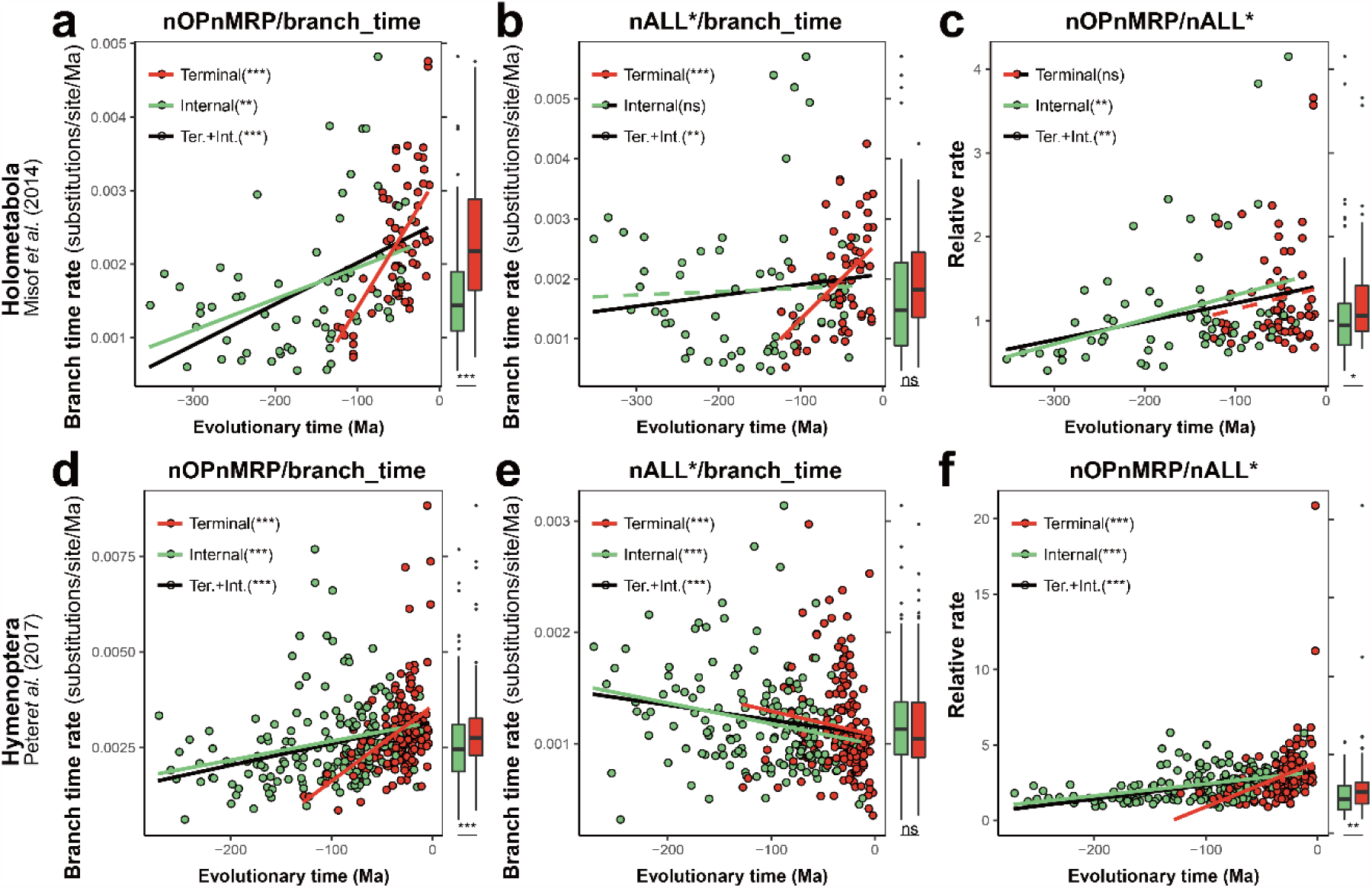
Mitochondria-associated nuclear-encoded proteins show different time-dependent evolutionary patterns compared to other nuclear-encoded proteins. **(a–c)** Correlations between evolutionary time and **(a)** branch time rates of combined nOP and nMRP (**nOPnMRP**), **(b)** nALL* and **(c)** relative rate of nOPnMRP, in Holometabola. **(d–f)** Same correlations are shown in Hymenoptera for the data set from Peters *et al.* 2017. Estimated dates at the midpoint of the branches were chosen for analysis. Red indicates terminal branches, and green indicates internal branches. Asterisks in parenthesis indicate significant level for non-parametric Spearman correlation; ns indicates *p* > 0.05; * *p* < 0.05; ** *p* < 0.01; *** *p* < 0.001.

Because haplodiploidy reduces the complicating factor of incomplete purifying selection on terminal branches, due to purging in the haploid sex (Werren 1993), we conducted additional analyses using a Hymenopteran data set (Peters *et al.*, 2017). The results show apparent rate acceleration over evolutionary time for mitochondria-associated nuclear proteins, but apparent rate deceleration for other nuclear proteins (Figure 5). However, the statistical significance of these results disappear following phylogenetic correction using a Bayesian mixed model (Table S5–6) (Hadfield & Nakagawa 2010). We therefore conclude that there is no clear evidence supporting evolutionary rate acceleration across time for mitochondria-associated nuclear-encoded proteins.

### 4. Using Evolutionary Rate Correlation to Predict Mitochondrial Interacting Nuclear Proteins

We next decided to investigate whether ERC can be used to predict candidate nuclear proteins previously unknown to interact with mitochondrial components. Here we focused on relative rate correlations with mOP, as mrRNA are more likely to be saturated. As a check on the predictive power of the method, we first tested whether ERC was a good predictor of known mitochondrial interacting nuclear proteins (i.e. nOP proteins in contact with mOP and nMRP proteins). After correction for multiple comparisons (FDR <0.05), 107 proteins were found with significant ERCs. Among these, 33 of 36 contacting nOP and nMRP proteins (91.7 %) were found to have significant ERC. For an FDR of 10%, 35/36 (97.2%) were significant, out of a set of 145 proteins with significant ERC proteins. Results therefore show that ERC is an effective predictor of proteins with known mitochondrial interactions.

Surprisingly, the phylogenetic contrasts ERC shows less power in detecting the known mOP contacted nOP and nMRP proteins (Figure S12), while at the same time reducing the number of significant correlations; only 11 of 36 (30.6%) contacting nOP and nMRP proteins were detected out of a total 20 proteins with significant ERC (FDR < 0.05), and the numbers are not much improved with a 10% FDR to 14/36 (38.9%) out of 26 proteins with significant ERC. We speculate that this reduced power may be due to noise introduced by the statistical methods for phylogenetic contrasts. Alternatively synergistic positive feedback between mitochondrial evolutionary rate and rates of substitution in interacting proteins (Rand 2004, Oliveira et al 2008) may result in stronger signals in those taxa with elevated mitochondrial mutation rates, with this signal being reduced following phylogenetic contrasts correction. We use the non-phylogenetically corrected ERCs to identify new candidate mitochondrial interacting proteins, but also report those found by the phylogenetic contrasts ERC (Table 2, S6).

**Table 2.**
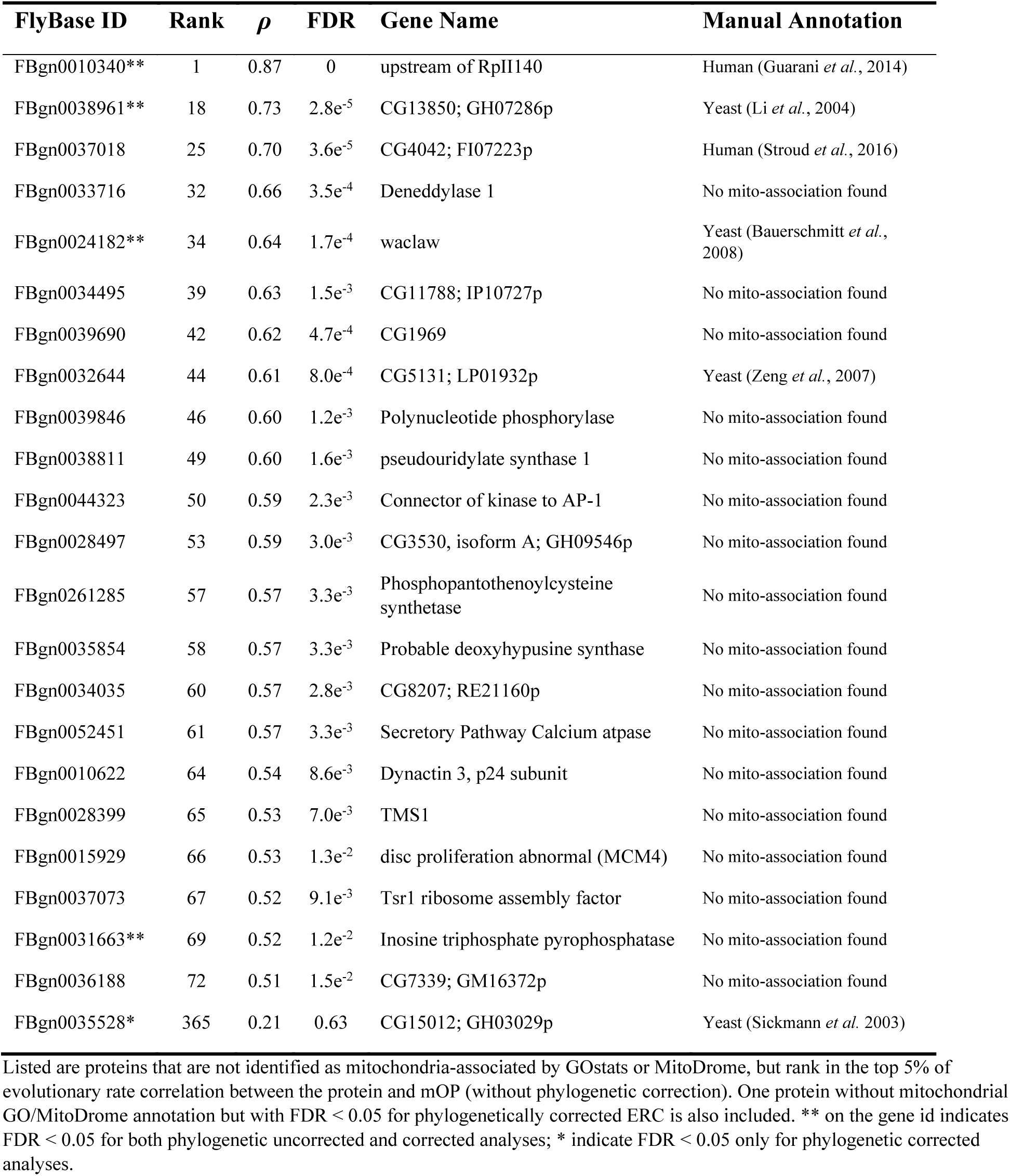
Mitochondria-associated nuclear protein candidates predicted in insects.

Approximately 7% of the single copy proteins (107 of 1478) show significant ERC with 5% FDR, with 60 proteins overlapping between this set and the138 mitochondria-associated proteins identified by GO/MitoDrome annotation (Figure S13). Thus, ERC at 5% FDR cut-off detects a significant number (47) of candidate interacting proteins not detected by standard gene ontology terms. Strikingly, the protein with the highest ERC (*ρ* = 0.87), known as “upstream of Rp1140” does not have a mitochondrial GO annotation. However, when we manually searched Google Scholar for relevant articles on this protein, a single article (Guarani *et al.*, 2014) was found implicating the putative human ortholog as a mitochondrial complex I assembly factor. We next conducted manual annotation by literature search of the proteins with the top 2% of ERC values (30 proteins of which 27 had prior mitochondrial gene ontology). Two additional proteins were uncovered without mitochondrial gene ontology but with papers implicating their putative orthologs as having mitochondrial function. Orthologs of FBgn0038961 and FBgn0037018, have been implicated to have mitochondria-associated function in yeast and humans (Li *et al.*, 2004; Stroud *et al.*, 2016), respectively. Thus, all 30 proteins (100%) with the highest ERC are implicated as having mitochondrial function, either by gene ontology (27) or annotation by our manual search (3), furtherly establishing the power of ERC for detecting nuclear proteins with mitochondrial function. Furthermore, the finding of the elevated ERCs in insects for the three proteins without prior mitochondrial gene ontology annotation, suggests that these proteins likely retain conserved mitochondrial functions beyond the taxa (human, yeast) in which their mitochondrial roles have so far been found.

Table 2 presents the set of 23 candidate genes without mitochondrial GO/MitoDrome annotation but which rank in the top 5% of 1478 phylogenetic uncorrected ERCs. The FDR for all are below 0.05. The list includes one additional protein (FBgn0035528) detected following phylogenetic correction but not in the phylogenetic uncorrected set, and four detected by both methods. Following manual annotation by literature survey, six were detected with some evidence of mitochondrial function (three described above). In addition, FBgn0035528 is a vacuolar membrane protein of unknown function in *Drosophila*, and its ortholog in yeast has been detected in the mitochondria (Sickmann *et al.*, 2003). Two additional genes are cited as showing mitochondrial associations in yeast (Bauerschmitt et al 2008, Zeng et al 2007), again supporting ERC as a means for detecting mitochondrial-interacting proteins and suggesting a conserved function also in insects. Remaining are 17 genes for which we have detected no literature supporting mitochondrial function, but which have elevated ERCs. Based on the ERC results, these are reasonable new candidates for mitochondrial interactions. It should be noted that there are different kinds of interaction between nuclear proteins and mitochondrial components that could result in elevated ERC. These could range from proteins coevolving in direct contact with mOP or the mitochondrial ribosome, to interactions within the mitochondrion, to downstream interactions in the cytosol.

GO enrichment analysis was conducted for 107 proteins with significant ERCs with FDR < 0.05. Almost all overrepresented “Cellular Component” GO items were contributed by mitochondria-related items, except minichromosome maintenance helicase (MCM) complex (Table S7). The MCM complex is involved in DNA replication, and is a heterohexameric complex comprised by 6 proteins (MCM2-7). Four proteins in the MCM complex are listed in the 1478 set and all are components of the cell cycle proteins (nCC), MCM 3, 4, 5 and 7. All four rank in the top 10% of 1478 proteins based on ERC. Two of the four MCM complex proteins were identified with FDR < 0.05, the other two were also identified with FDR < 0.10 (Table S3). These results suggest a previously unknown interaction between the MCM complex and mitochondria that result in a correlation in the evolutionary rates.

These results indicate that ERC is a powerful tool for detecting nuclear proteins that interact with mitochondrial products over evolution, and has potential to detect new candidate nuclear proteins and protein complexes not previously known to have mitochondrial interactions or functions.

## Conclusions

Our study provides evidence for strong evolutionary rate coevolution between mitochondrial products and mitochondria-associated nuclear-encoded proteins in insects. Physical contacts with mitochondrial products show even stronger rate correlation. Our study also revealed different patterns across evolutionary time of nuclear proteins that interact with mitochondria compared to other nuclear-encoded proteins. Evolutionary rate correlation (ERC) of nuclear and mitochondrial proteins is a strong predictor of mitochondria interacting proteins, and shows promise as a method for detecting previously unknown mitochondria interacting proteins.

## Methods

### Retrieving mitochondrial genomes

For each 64 holometabolous species in Misof *et al.* (2014) paper, we searched available mitochondrial genomes from either the same insect species or a relative. Using the Cox1 sequence from each species, online blastn was conducted by restricting “organism” to the same family and setting “max target sequences” as 20000 (2017-Nov). If the Cox1 sequence of that species was not available, Cox1 sequence of the closest taxon was retrieved, and used to retrieve the mitochondrial sequences of the related species. By this method, 25 mitochondrial genomes were retrieved from the same species or genus, 7 from the same subfamily and 11 from the same family. An additional mitochondrial genome was also included from *Hydropsyche pellucidula*, whose corresponding species in Misof *et al.* (2014) tree is an uncharacterized specie in the same suborder Annulipalpia (Table S1), resulting in 44 holometabolous mitochondrial genomes. Five hemipterans were used as outgroups. GenBank accessions of mitochondrial genomes were listed in Table S1.

### Phylogenetic analysis for mitochondrial proteins and rRNA

mOP protein and mrRNA sequences were extracted from GenBank files, except for *Leptopilina boulardi* mitochondrial genome which was annotated using MITOS webserver. Three degenerate bases W and 2 R were converted into A for 12S rRNA in *Jellisonia amadoi*. mOP proteins were aligned using Mafft v7.123b with default parameters. mrRNAs were aligned using online R-coffee (Moretti *et al.*, 2008) to account for secondary structure. Alignments were filtered by trimAI version 1.2rev59 with automated settings (Capella-Gutierrez *et al.*, 2009), then concatenated using AMAS (Borowiec 2016). Based on Misof *et al.* (2014) tree topology, branch lengths of mOP and mrRNA were estimated using RAxML v8.0.20, applying MTART+G+I and GTR+G+I model, respectively. For concatenated mOP and mrRNA, best substitution models were determined using ProtTest v3.4.2 (Darriba *et al.*, 2011) and jModelTest v2.1.10 (*Darriba et al., 2012*), respectively. “+F” was not considered for model selection, as “-t” option in RAxML doesn’t support “+F”. Branch lengths and corresponding branch times were extracted using Newick Utilities v1.6 (Junier & Zdobnov 2010).

### Defining nuclear-encoded protein categories

For defining nuclear-encoded protein categories, KEGG or GO annotations were determined based on their *Drosophila* orthologs in FlyBase v2017_06 (Gramates *et al.*, 2017). The GO annotations of *Drosophila* proteins were retrieved directly from FlyBase v2017_06. The KEGG annotation of *Drosophila* proteins were annotated using the BlastKOALA online service (Kanehisa *et al.*, 2016).

To identify nuclear proteins that are mitochondria-associated, we retrieved genes with or belong to the cellular component term “mitochondrion” (GO: 0005739), using R package GOstats v2.36.0 (Falcon & Gentleman 2007). Of the 1478 single copy genes in our analysis, 132 were identified as mitochondria-associated. Six additional proteins were identified using information from MitoDrome database, for a total of 138 proteins annotated by GOstats or MitoDrome database as mitochondria-associated.

Among the 138 mitochondria-associated nuclear proteins, we next undertook to identify nuclear-encoded OXPHOS proteins and mitochondrial ribosomal proteins. Nineteen of 1478 single copy genes were identified with KEGG annotation as “oxidative phosphorylation (OP)” (map00190). Three of 19 were vacuolar-type ATPase proteins and filtered out, as vacuolar-type ATPase are found within the membranes of many organelles. Finally, 16 were defined as nuclear-encoded oxidative phosphorylation proteins (nOP). 27 were identified with GO annotation “mitochondrial ribosome” (GO:0005761), “mitochondrial large ribosomal subunit” (GO:0005762) or “mitochondrial small ribosomal subunit” (GO:0005763). One (EOG5DV42G) of 27 was a 5-methylcytosine rRNA methyltransferase and manually filtered out; 26 were defined as nuclear-encoded mitochondrial ribosomal proteins (nMRP). The 96 remaining mitochondria-associated proteins, minus those identified as nuclear-encoded OXPHOS or mitochondrial ribosomal proteins, were designated as other mitochondria-associated nuclear proteins (OMito), and are analyzed separately in some treatments.

To address the contribution of physical contact to evolutionary rate correlation, nOP proteins were furtherly separated into two categories based on their complex structure information (Jonckheere *et al.*, 2012; Kuhlbrandt 2015; Tsukihara *et al.*, 1996; Yoshikawa *et al.*, 2012; Zhu *et al.*, 2016). One category is 10 nOP proteins in contact with mOPs, and the other is 4 nOP proteins not in contact with mOPs. Two proteins whose mOP contacting information couldn’t be determined were excluded. They are cytochrome c oxidase assembly protein COX15-like protein (EOG563XTD) and cytochrome c oxidase assembly protein COX11 (EOG5RBP1H).

Based on structure of the mammalian (*Sus scrofa*) mitochondrial ribosomal comple*x (Greber et al., 2015*; Greber *et al.*, 2014), all nMRPs directly contact with mrRNAs, except mRpL1 (EOG508KQM) which is not present in the structure. Amino acid sites were separated into sites directly in contact with mitochondrial rRNA/tRNA and those not. The contacts between nMRPs and mitochondrial rRNAs/tRNAs were determined using “InterfaceResidues” function in PyMOL v1.7.0.0 (http://pymol.sourceforge.net/). *Sus scrofa* mitochondrial ribosomal proteins were aligned to corresponding original alignments using Mafft v7.123b (Katoh & Standley 2013) with “-add” option. As not all amino acids have been solved in the complex, only the solved amino acids were extracted based on their structure information. The alignments were further filtered based on provided Aliscore results from Misof *et al.* (2014) paper.

The remaining 1340 nuclear proteins (minus nOP, nMRP, and OMito) are designated as nALL*. Among these, we identified two control categories for comparative purposes. Nineteen proteins were identified with the KEGG annotation “Cell Cycle” (map04110), and defined as nuclear-encoded cell cycle proteins (nCC). Thirty-five proteins were identified with GO annotation “cytosolic large ribosomal subunit” (GO:0022625), “cytosolic ribosome” (GO:0022626) or “cytosolic small ribosomal subunit” (GO:0022627). Twenty-nine of these 35 were finally defined as nuclear-encoded cytosolic ribosomal proteins (nCRP) after manually curation. These cytosolic ribosomal proteins serve as a useful control for comparisons to nuclear-encoded mitochondrial ribosomal proteins.

### Phylogenetic analysis for nuclear protein categories

For evolutionary rate estimation, protein alignments of the 1478 single copy nuclear genes and the dated tree were retrieved from supplementary materials of Misof *et al.* (2014). Both original and filtered alignments, which were filtered by Aliscore (Misof *et al.*, 2014), were retrieved and subjected to analysis. Although branch lengths estimated using filtered alignments are much shorter than those using original alignments, we found that the resulting patterns and conclusions were quite similar. Here we only present results generated using filtered alignments.

For nuclear categories, the alignments were concatenated using AMAS (Borowiec 2016). Branch lengths were estimated using RAxML v8.0.20 (Stamatakis 2014) with the Misof *et al.* (2014) tree topology as the given input tree. The substitution model used is “LG+G+I”, which was determined using ProtTest v3.4.2 (Darriba *et al.*, 2011). Model “LG+G” was also tested, and the results were almost the same. When compared to the mitochondrial evolutionary rates and distances to HCA, the nuclear trees and mitochondrial trees were pruned to the same corresponding topology using Newick Utilities v1.6 (Junier & Zdobnov 2010). Branch lengths and corresponding branch times were extracted using Newick Utilities v1.6 (Junier & Zdobnov 2010). The trees were visualized using iTOL (Letunic & Bork 2016).

### Phylogenetic comparative analysis

For phylogenetic correction of traits on terminal branches, non-parametric correlation analysis were conducted for phylogenetic independent contrasts. Independent contrasts were extracted using R package Ape v4.1. Spearman correlations through origins were calculated using R package Picante v1.6.2.

As conventional phylogenetic correction is not applied to traits on internal branches, we used Bayesian phylogenetic mixed models to test hypotheses about the correlation of evolutionary rates (internal and terminal) and evolutionary time. The response variable in these analyses is evolutionary rates. Evolutionary times were explored as explanatory variables, and treated as fixed effects in the model. Estimated time from midpoint of branches to present were used as evolutionary time. Misof *et al.* (2014) phylogeny was included in the analysis as a random effect to control for correlations due to shared evolutionary history. The MCMC was run for five million iterations with a thinning interval of 500 and a ‘burn in’ of 1000. Convergence of the chains was confirmed by visual inspection of the trace plots. For the fixed effect, we used a uniform prior (Hadfield & Nakagawa 2010). All generalized linear mixed models were implemented using R package MCMCglmm v.2.24 (Hadfield & Nakagawa 2010).

### Individual nuclear gene analysis

For each nuclear protein, branch length was estimated based on the Misof *et al.* (2014) tree using RAxML v8.0.20 by setting substitution model as “PROTGAMMAIAUTO”. As not all nuclear individual alignments cover 44 selected holometabolous species, both constructed mitochondrial and nuclear trees were pruned to the same corresponding topology for mitochondrial-nuclear correlation analysis. The coverage number of 44 selected holometabolous species were provided in supplementary Table S3. Branch lengths and corresponding branch times were extracted using Newick Utilities v1.6 (Junier & Zdobnov 2010) for each pruned tree. To treat every protein equally, the concatenated set of all 1478 genes was used for relative rate normalization of individual proteins. nALL* was also tested for relative rate normalization, showing similar results.

### GO enrichment analysis

GO enrichment analysis was conducted using R package GOstats v2.36.0 (Falcon & Gentleman 2007), applying the conditional hypergeometric test. Since GO items are often members of the same hierarchy, tests are dependent and not suitable for direct multiple correction. As recommended by manual of GOstats, returned conditional *p* values were interpreted as corrected and directly used.

### Subsampling analysis

To test potential effects of special features of mitochondria-associated nuclear proteins, we employed subsampling methods (Politis *et al.*, 1999). Five subsampling comparisons were made to control for gene number, evolutionary rate, alignment length, conservation score and gap percentage, respectively. For each method, 1000 replicates were randomly generated. In every replicate, for controlling gene number, genes were randomly sampled from nALL* category to match up the gene number in treatment gene category. For controlling evolutionary rate, genes were randomly sampled from the nALL* category with matched evolutionary rates to genes in treatment gene category. Here evolutionary rate for each protein was defined and calculated as the sum of internal and terminal branch lengths in the Holometabola divided by the sum of their corresponding branch times. Evolutionary rates of nOP and nMRP proteins range from 0.00057 to 0.0038 aa/sites/Mya. Totally 1167 proteins have evolutionary rates range from 0.0005 to 0.0040 aa/sites/Mya. These 1167 proteins plus nOP and nMRP proteins were divided into 100 bins. Proteins in the same bin were treated as proteins with matched evolutionary rates. For controlling alignment length, blocks were randomly sampled from concatenated nALL* alignments with matched alignment lengths to genes in treatment gene category. For controlling gap percentage in 64 holometabolous species, amino acid sites were randomly sampled from concatenated nALL* alignments with matched gap percentage to sites in concatenated treatment alignment. Here undetermined characters were also treated as gaps, because RAxML treats gap and undetermined characters technically equal. For controlling conservation score, amino acid sites were randomly sampled from concatenated nALL* alignments with matched conservation scores to sites in concatenated treatment alignment. Conservation scores for amino acid sites were calculated using Protein Residue Conservation Prediction online service http://compbio.cs.princeton.edu/conservation/score.html (Capra & Singh 2007). The method used is Jensen-Shannon divergence score. The window size was set as 1. The background and matrix were both set as BLOSUM62. Conservation scores ranges from 0 to 1, and were formatted into 2 digits after the decimal point for subsampling convenience. All sampling described in this section are without replacement for each replicate. Sampled genes, blocks or sites were concatenated using AMAS (Borowiec 2016). Branch lengths were estimated using RAxML v8.0.20 with option “-t” and substitution model LG+G+I, and extracted using Newick Utilities v1.6 (Junier & Zdobnov 2010).

### Statistical analysis

Correlation analyses were conducted using the two-sided Spearman method. Comparisons of dependent Spearman correlations without deattentuation were performed using methods and scripts from Rosner *et al.* (2015) in SAS v9.4b. Statistical tests among samples were conducted using the Kruskal–Wallis test. Post-hoc tests were performed by Fisher’s least significant difference using the function “kruskal” in the R package Agricolae v1.2.6 by setting the multiple correction method as “fdr”. Pairwise comparison between Hymenoptera and Diptera were performed using a two-sided Wilcoxon rank sum test. The Benjamini & Hochberg method was used for multiple corrections. Statistical analyses were performed in R v3.2.0.

## Acknowledgements

This research was supported by grants from the US National Science Foundation (DEB-1257053 and IOS-1456233) and Nathaniel & Helen Wisch Chair to JHW, Major International (Regional) Joint Research Project of NSFC (Grant no. 31620103915 to GY and JHW), National Natural Science Foundation of China (Grant no. 31472038 to GY, no. 31701843 to ZY) and China Postdoctoral Science Foundation (2016M601947) to ZY. We thank Dr. Xin Zhou (China Agricultural University) for providing a dated insect phylogenetic tree file. We thank Dr. Martin Kaltenpoth (Johannes Gutenberg University Mainz) for providing some insect mitochondrial sequences, and Yu Tian and Jiang Hu (Nextomics Biosciences Co., Ltd.) for helping with phylogenetic analyses. We thank John Jaenike, Rachel Edwards and Philip Bellomio (University of Rochester, USA) for their comments and suggestions on the manuscript, and Xinhai Ye (Zhejiang University, China) for refining figures.

## Author Contributions

JHW and ZY conceived and designed the research; GY supervised the work of ZY and provided feedback during development of the project; ZY performed analyses; JHW and ZY interpreted the results; JHW, ZY and GY wrote and revised the manuscript.

