## Supplementary Materials for "Evolutionary rate coevolution between mitochondria and mitochondria-associated nuclear-encoded proteins in insects"

**This supplemental file includes**

Table S1–2, S4–8

Figure S1 to S27

Supplementary text

Supplementary methods

References

**Other supplementary material for this manuscript includes the following:**

Table S3 (provided as separate excel file)

#### Supplementary tables

**Table S1.** Taxonomy and GenBank accessions of mitochondrial genomes used in this paper.

**Table S2.** *P* values for correlations between mitochondrial products and mitochondria-associated nuclear-encoded proteins after considering order effect.

**Table S3.** Correlation of individual nuclear-encoded protein and mOP relative rates. (Provided as separate excel file)

**Table S4.** Comparison of evolutionary rate correlation with mitochondrial products among nuclear protein categories using individual gene analyses.

**Table S5.** Phylogenetic corrected *P* values for correlations between evolutionary rates and evolutionary time in Holometabola.

**Table S6.** Phylogenetic corrected *P* values for correlations between evolutionary rates and evolutionary time in Hymenoptera.

**Table S7.** Overrepresented GO "Cellular Component" terms for proteins which are significantly correlated with mOP relative rate (FDR < 0.05).

**Table S8.** Phylogenetic corrected *P* values for correlations between evolutionary rates and evolutionary time in Holometabola excluding Hymenoptera clade.

#### Supplementary figures

**Figure S1.** Correlations between distances to HCA of mitochondrial products and nuclear encoded protein categories.

**Figure S2.** Correlations between branch time (BTime) rates of mitochondrial products and nuclear encoded protein categories.

**Figure S3.** Correlations between standardized contrasts of relative evolutionary rates of mitochondrial products and nuclear encoded protein categories.

**Figure S4.** Correlations between standardized contrasts of branch time evolutionary rates of mitochondrial products and nuclear encoded protein categories.

**Figure S5.** Subsampling statistics for (a) nOP-mOP branch time (BTime) evolutionary rate correlation (ERC) and (b) nMRP-mrRNA ERC.

**Figure S6.** Amino acid composition of different concatenated nuclear-encoded protein categories.

**Figure S7.** Mitochondria-associated nuclear-encoded proteins in contact with mitochondrial products show stronger branch time (BTime) evolutionary rate correlations (ERCs) with mitochondrial products.

**Figure S8.** Structure of mammalian mitochondrial ribosome.

**Figure S9.** Correlations between standardized contrasts of relative evolutionary rates of mitochondrial products and nuclear encoded protein categories in contact or not in contact with mitochondrial products.

**Figure S10.** Correlations between standardized contrasts of branch time rates of mitochondrial products and nuclear encoded protein categories in contact or not in contact with mitochondrial products.

**Figure S11.** Subsampling statistics for correlations between evolutionary rates of nOPnMRP and their evolutionary time.

**Figure S12.** Correlations of nuclear and mOP relative rates before and after phylogenetic correction.

**Figure S13.** Venn graph of different nuclear-encoded protein categories.

**Figure S14.** Relative evolutionary rates of concatenated mitochondrial products and nuclear protein categories vary across taxa in Holometabola.

**Figure S15.** Dated phylogenetic trees in Holometabola.

**Figure S16.** Branch time (BTime) evolutionary rates of concatenated mitochondrial products and nuclear protein categories vary across insect orders in Holometabola.

**Figure S17.** Evolutionary rate ratios of components in contact over those not in contact with mitochondrial products in Holometabola.

**Figure S18.** Relative rates of mitochondria-associated nuclear-encoded proteins show acceleration within Psocodea.

**Figure S19.** Relative rate variation of mitochondria-associated nuclear-encoded proteins in Psocodea.

**Figure S20.** Correlations between branch time (BTime) evolutionary rates and their estimated evolutionary time in Holometabola.

**Figure S21.** Subsampling statistics for correlations between branch time (BTime) evolutionary rates and their evolutionary time (ETime).

**Figure S22.** Correlations between evolutionary rates and their estimated evolutionary time in Holometabola excluding Hymenoptera clade.

**Figure S23.** Correlations between branch time (BTime) evolutionary rates and their estimated evolutionary time in Hymenoptera.

**Figure S24.** Correlations between relative evolutionary rates and their estimated evolutionary time in Holometabola.

**Figure S25.** Subsampling statistics for correlations between relative evolutionary rates and their evolutionary time (ETime).

**Figure S26.** Correlations between relative evolutionary rates and their estimated evolutionary time in Hymenoptera.

**Figure S27.** Relation of evolutionary rate correlation (ERC) with mOP and apparent acceleration across evolutionary time (ETime).

#### Supplementary text

**S1.** Further analyses on evolutionary patterns of mitochondrial products and mitochondria-associated nuclear-encoded proteins in Holometabola

**S2.** Mitochondrial-nuclear evolutionary rate coevolution in Psocodea

**S3.** Apparent acceleration of nuclear proteins associated with mitochondria

**S3.1** Correlation between branch time rates and evolutionary time

**S3.2** Correlation between relative evolutionary rates and evolutionary time

**S3.3** Individual gene analysis for apparent acceleration of mitochondria- associated nuclear-encoded proteins

**Table S1. Taxonomy and GenBank accessions of mitochondrial genomes used in this paper.**

| Species | Order | Family | Corresponding species in Misof *et al.* tree | Shared taxon level | Accession ID |
|---|---|---|---|---|---|
| ***Nilaparvata lugens*** | Hemiptera | Delphacidae | same | specie | NC_021748.1 |
| ***Acyrthosiphon pisum*** | Hemiptera | Aphididae | same | specie | NC_011594.1 |
| ***Aphis gossypii*** | Hemiptera | Aphididae | same | specie | NC_024581.1 |
| ***Trialeurodes vaporariorum*** | Hemiptera | Aleyrodidae | same | specie | NC_006280.1 |
| ***Bemisia tabaci*** | Hemiptera | Aleyrodidae | same | specie | NC_006279.1 |
| ***Apis mellifera*** | Hymenoptera | Apidae | same | specie | NC_001566.1 |
| ***Bombus terrestris*** | Hymenoptera | Apidae | same | specie | KT368150.1 |
| ***Orussus occidentalis*** | Hymenoptera | Orussidae | *Orussus abietinus* | genus | FJ478174.1 |
| ***Cotesia vestalis*** | Hymenoptera | Braconidae | same | specie | NC_014272.1 |
| ***Tenthredo tienmushana*** | Hymenoptera | Tenthredinidae | *Tenthredo koehleri* | genus | KR703581.1 |
| ***Leptopilina boulardi*** | Hymenoptera | Figitidae | *Leptopilina clavipes* | genus | KU665622.1 |
| ***Nasonia vitripennis*** | Hymenoptera | Pteromalidae | same | specie | EU746609.1<br>EU746613.1 |
| ***Atta texana*** | Hymenoptera | Formicidae | *Acromyrmex echinatior* | tribe | MF417380.1 |
| ***Mylabris* sp.** | Coleoptera | Meloidae | *Meloe violaceus* | family | JX412732.1 |
| ***Aleochara* sp.** | Coleoptera | Staphylinidae | *Aleochara curtula* | genus | KT780622.1 |
| ***Macrogyrus oblongus*** | Coleoptera | Gyrinidae | *Gyrinus marinus* | family | FJ859901.1 |

| Species | Order | Family | Corresponding species in Misof *et al.* tree | Shared taxon level | Accession ID |
|---|---|---|---|---|---|
| ***Damaster mirabilissimus*** | Coleoptera | Carabidae | *Carabus granulatus* | genus (a) | GQ344500.1 |
| ***Tribolium castaneum*** | Coleoptera | Tenebrionidae | same | specie | NC_003081.2 |
| ***Tomicus piniperda*** | Coleoptera | Curculionidae | *Dendroctonus ponderosae* | subfamily | KX035226.1 |
| ***Mengenilla moldrzyki*** | Strepsiptera | Mengenillidae | same | specie | NC_018545.1 |
| ***Sialis* sp. BMNH 1425199** | Megaloptera | Sialidae | *Sialis lutaria* | genus | KT876911.1 |
| ***Corydalus cornutus*** | Megaloptera | Corydalidae | same | specie | NC_011276.1 |
| ***Myrmeleon immanis*** | Neuroptera | Myrmeleontidae | *Euroleon nostras* | subfamily | KJ461323.1 |
| ***Thyridosmylus langii*** | Neuroptera | Osmylidae | *Osmylus fulvicephalus* | family | KC515397.1 |
| ***Semidalis aleyrodiformis*** | Neuroptera | Coniopterygidae | *Conwentzia psociformis* | subfamily | KT425067.1 |
| ***Mongoloraphidia harmandi*** | Raphidioptera | Raphidiidae | *Xanthostigma xanthostigma* | famliy | FJ859902.1 |
| ***Anabolia bimaculata*** | Trichoptera | Limnephilidae | *Platycentropus radiatus* | tribe | MF680449.1 |
| ***Hydropsyche pellucidula*** | Trichoptera | Hydropsychidae | *Annulipalpia* sp. AD-2013 | suborder (b) | NC_029246.1 |
| ***Prays oleae*** | Lepidoptera | Yponomeutidae | *Yponomeuta evonymella* | family | KM874804.1 |
| ***Napialus hunanensis*** | Lepidoptera | Hepialidae | *Triodia sylvina* | family | KJ632465.1 |
| ***Manduca sexta*** | Lepidoptera | Sphingidae | same | specie | NC_010266.1 |

| Species | Order | Family | Corresponding species in Misof *et al.* tree | Shared taxon level | Accession ID |
|---|---|---|---|---|---|
| ***Parasa consocia*** | Lepidoptera | Zygaenidae | *Zygaena fausta* | family | KX108765.1 |
| ***Bombyx mori*** | Lepidoptera | Bombycidae | same | specie | NC_002355.1 |
| ***Celastrina hersilia*** | Lepidoptera | Lycaenidae | *Polyommatus icarus* | subfamily | HM243589.1 |
| ***Pachliopta aristolochiae*** | Lepidoptera | Papilionidae | *Parides eurimedes* | tribe | KU950357.1 |
| ***Drosophila melanogaster*** | Diptera | Drosophilidae | same | specie | NC_024511.2 |
| ***Exorista sorbillans*** | Diptera | Tachinidae | *Triarthria setipennis* | family | HQ322500.1 |
| ***Sarcophaga crassipalpis*** | Diptera | Sarcophagidae | same | specie | NC_026667.1 |
| ***Ceratitis capitata*** | Diptera | Tephritidae | *Rhagoletis pomonella* | family | AJ242872.1 |
| ***Aedes aegypti*** | Diptera | Culicidae | same | specie | NC_035159.1 |
| ***Anopheles gambiae*** | Diptera | Culicidae | same | specie | NC_002084.1 |
| ***Phlebotomus papatasi*** | Diptera | Psychodidae | same | specie | NC_028042.1 |
| ***Trichocera bimacula*** | Diptera | Trichoceridae | *Trichocera saltator* | genus | NC_016169.1 |
| ***Tipula cockerelliana*** | Diptera | Tipulidae | *Tipula maxima* | genus | NC_030520.1 |
| ***Jellisonia amadoi*** | Siphonaptera | Ceratophyllidae | *Ceratophyllus gallinae* | family | KF322091.1 |
| ***Nannochorista philpotti*** | Mecoptera | Nannochoristidae | same | specie | HQ696580.1 |
| ***Neopanorpa pulchra*** | Mecoptera | Panorpidae | *Panorpa vulgaris* | family | JX569848.1 |

| Species | Order | Family | Corresponding species in Misof *et al.* tree | Shared taxon level | Accession ID |
|---|---|---|---|---|---|
| ***Boreus elegans*** | Mecoptera | Boreidae | *Boreus hyemalis* | genus | NC_015119.1 |
| ***Bittacus pilicornis*** | Mecoptera | Bittacidae | same | specie | NC_015118.1 |

(a) *Damaster* is subgenus in genus *Carabus*.

(b) Corresponding specie in Misof *et al.* tree is an unclassified species in suborder Annulipalpia.

**Table S2. *P* values for correlations between mitochondrial products and mitochondria-associated nuclear-encoded proteins after considering order effect.** *P* values were calculated for linear regressions using insect order as variable and Spearman correlations excluding hymenopterans. Significant correlations ($p < 0.05$) are indicated by *.

| Correlation | Relative rate | | Branch time rate | |
|---|---|---|---|---|
| | ***p*** (order as variable) | ***p*** (exclude Hym) | ***p*** (order as variable) | ***p*** (exclude Hym) |
| mOP-nOP | *2.0e$^{-9}$ | *3.0e$^{-4}$ | *7.3e$^{-8}$ | *1.4e$^{-4}$ |
| mOP-nALL* | NA | NA | 0.16 | *1.9e$^{-3}$ |
| mOP-nCC | 0.98 | 0.86 | 0.12 | *8.7e$^{-3}$ |
| mrRNA-nMRP | *1.9e$^{-11}$ | *6.5e$^{-5}$ | *3.2e$^{-10}$ | *2.5e$^{-6}$ |
| mrRNA-nALL* | NA | NA | 0.14 | *1.4e$^{-3}$ |
| mrRNA-nCRP | 0.56 | 0.52 | 0.63 | 0.19 |

**Table S3. Correlation of individual nuclear-encoded protein and mOP relative rates.** (Provided as separate excel file)

**Table S4. Comparison of evolutionary rate correlation with mitochondrial products among nuclear protein categories using individual gene analyses.** Pairwise Wilcoxon rank sum tests were conducted among ERCs between nuclear-encoded categories and mitochondrial products. *P* values were adjusted using the Benjamini & Hochberg method for multiple correction. Comparisons on the left in the first column have higher correlation values than those on the right.

| Correlation ($\rho$) comparison (Wilcoxon rank sum test) | | | Relative rate | | | Branch time rate | | |
|---|---|---|---|---|---|---|---|---|
| | | | **W** | ***p*** | **adjusted *p*** | **W** | ***p*** | **adjusted *p*** |
| nOP-mOP | vs. | nCC-mOP | 266 | $6.2e^{-5}$ | $7.5e^{-5}$ | 255 | $3.8e^{-4}$ | $4.6e^{-4}$ |
| nOP-mOP | vs. | nALL*-mOP | 18517 | $5.1e^{-8}$ | $7.7e^{-8}$ | 17404 | $2.5e^{-6}$ | $3.8e^{-6}$ |
| nOPcon-mOP | vs. | nOPnot-mOP | 40 | $2.0e^{-3}$ | $2.0e^{-3}$ | 40 | $2.0e^{-3}$ | $2.0e^{-3}$ |
| nMRP-mrRNA | vs. | nCRP-mrRNA | 754 | $5.6e^{-16}$ | $1.1e^{-15}$ | 754 | $5.6e^{-16}$ | $1.7e^{-15}$ |
| nMRP-mrRNA | vs. | nALL*-mrRNA | 33375 | $6.3e^{-18}$ | $1.9e^{-17}$ | 33342 | $7.3e^{-18}$ | $4.4e^{-17}$ |
| OMito-mOP | vs. | nALL*-mOP | 96535.5 | $8.2e^{-20}$ | $4.9e^{-19}$ | 91715.5 | $4.6e^{-15}$ | $9.1e^{-15}$ |
| OMito-mrRNA | vs. | nALL*-mrRNA | 100032 | $1.1e^{-23}$ | $7.6e^{-23}$ | 97552 | $6.7e^{-21}$ | $4.7e^{-20}$ |

**Table S5. Phylogenetic corrected *P* values for correlations between evolutionary rates and evolutionary time in Holometabola.** Bayesian mixed model was performed using MCMCglmm for phylogenetic correction. *P* values were adjusted using Benjamini & Hochberg method for multiple correction. Significant *P* values (< 0.05) for positive correlations between evolutionary rate and evolutionary time are indicated by *. BTime: branch time. Note that although *P* for nCC/nALL* is significant, but it shows negative correlation with evolutionary time.

| | **Int.+Ter.** | | **Terminal** | | **Internal** | |
|---|---|---|---|---|---|---|
| | ***p*** | **adjusted *p*** | **p** | **adjusted *p*** | ***p*** | **adjusted *p*** |
| nOP/nALL* | *0.002 | *0.009 | 0.18 | 0.81 | *0.0002 | *0.0018 |
| nMRP/nALL* | *0.0001 | *0.0009 | 0.14 | 0.81 | *0.0004 | *0.0018 |
| nCC/nALL* | 0.013 | 0.04 | 0.56 | 0.97 | 0.66 | 0.97 |
| nCRP/nALL* | 0.29 | 0.65 | 0.92 | 0.97 | 0.93 | 0.97 |
| nALL*/BTime | 0.85 | 0.99 | 0.93 | 0.97 | 0.94 | 0.97 |
| nOP/BTime | 0.99 | 0.99 | 0.97 | 0.97 | 0.96 | 0.97 |
| nMRP/BTime | 0.89 | 0.99 | 0.92 | 0.97 | 0.95 | 0.97 |
| nCC/BTime | 0.97 | 0.99 | 0.97 | 0.97 | 0.97 | 0.97 |
| nCRP/BTime | 0.98 | 0.99 | 0.96 | 0.97 | 0.96 | 0.97 |

**Table S6. Phylogenetic corrected *P* values for correlations between evolutionary rates and evolutionary time in Hymenoptera.** Data was retrieved from Peter *et al.* (2017). Bayesian mixed model was performed using MCMCglmm for phylogenetic correction. *P* values were adjusted using Benjamini & Hochberg method for multiple correction. Significant *P* values (< 0.05) for positive correlations between evolutionary rate and evolutionary time are indicated by *. BTime: branch time. Note that although *P* for nCC/nALL* is significant, but it shows negative correlation with evolutionary time.

| | **Int.+Ter.** | | **Terminal** | | **Internal** | |
|---|---|---|---|---|---|---|
| | ***p*** | **adjusted *p*** | ***p*** | **adjusted *p*** | ***p*** | **adjusted *p*** |
| nOP/nALL* | *0.002 | *0.009 | *0.003 | *0.014 | 0.17 | 0.51 |
| nMRP/nALL* | *0.0001 | *0.0009 | *0.0008 | *0.0072 | 0.061 | 0.51 |
| nCC/nALL* | 0.01 | 0.024 | 0.031 | 0.069 | 0.17 | 0.51 |
| nCRP/nALL* | *0.03 | 0.064 | *0.012 | *0.036 | 0.6 | 0.98 |
| nALL*/BTime | 0.94 | 0.96 | 1 | 1 | 0.95 | 0.98 |
| nOP/BTime | 0.73 | 0.96 | 0.73 | 1 | 0.94 | 0.98 |
| nMRP/BTime | 0.9 | 0.96 | 0.91 | 1 | 0.95 | 0.98 |
| nCC/BTime | 0.9 | 0.96 | 0.98 | 1 | 0.93 | 0.98 |
| nCRP/BTime | 0.96 | 0.96 | 0.94 | 1 | 0.98 | 0.98 |

**Table S7. Overrepresented GO "Cellular Component" terms for proteins which are significantly correlated with mOP relative rate (FDR < 0.05).**

| GO ID | Term | *P* value | Count | Size |
|---|---|---|---|---|
| GO:0000313 | organellar ribosome | $2.9e^{-28}$ | 25 | 27 |
| GO:0005759 | mitochondrial matrix | $1.0e^{-27}$ | 30 | 42 |
| GO:0005762 | mitochondrial large ribosomal subunit | $2.8e^{-16}$ | 15 | 17 |
| GO:0005763 | mitochondrial small ribosomal subunit | $2.3e^{-12}$ | 10 | 10 |
| GO:0005747 | mitochondrial respiratory chain complex I | $1.3e^{-10}$ | 10 | 12 |
| GO:0098803 | respiratory chain complex | $1.3e^{-10}$ | 10 | 12 |
| GO:0030964 | NADH dehydrogenase complex | $1.3e^{-10}$ | 10 | 12 |
| GO:0005743 | mitochondrial inner membrane | $2.1e^{-9}$ | 14 | 30 |
| GO:0005740 | mitochondrial envelope | $1.2e^{-8}$ | 14 | 34 |
| GO:0015934 | large ribosomal subunit | $1.4e^{-8}$ | 15 | 39 |
| GO:0098798 | mitochondrial protein complex | $2.9e^{-8}$ | 10 | 17 |
| GO:0043233 | organelle lumen | $9.6e^{-8}$ | 39 | 247 |
| GO:0044455 | mitochondrial membrane part | $1.2e^{-7}$ | 10 | 19 |
| GO:0015935 | small ribosomal subunit | $2.3e^{-7}$ | 10 | 20 |
| GO:0031975 | envelope | $7.2e^{-7}$ | 16 | 57 |
| GO:0005739 | mitochondrion | $9.0e^{-7}$ | 11 | 52 |
| GO:0030529 | intracellular ribonucleoprotein complex | $9.7e^{-6}$ | 28 | 174 |
| GO:0005737 | cytoplasm | $3.9e^{-5}$ | 72 | 753 |
| GO:0043227 | membrane-bounded organelle | $9.8e^{-5}$ | 59 | 680 |
| GO:0031090 | organelle membrane | $1.5e^{-3}$ | 17 | 111 |
| GO:0043232 | intracellular non-membrane-bounded organelle | $4.8e^{-3}$ | 31 | 281 |
| GO:0016507 | mitochondrial fatty acid beta-oxidation multienzyme complex | $5.1e^{-3}$ | 2 | 2 |
| GO:0032991 | macromolecular complex | $6.2e^{-3}$ | 53 | 570 |
| GO:0043229 | intracellular organelle | $6.6e^{-3}$ | 46 | 598 |
| GO:0005758 | mitochondrial intermembrane space | 0.015 | 2 | 3 |

| GO ID | Term | *P* value | Count | Size |
|---|---|---|---|---|
| GO:0042555 | MCM complex | 0.028 | 2 | 4 |
| GO:0044429 | mitochondrial part | 0.037 | 1 | 1 |

**Table S8. Phylogenetic corrected *P* values for correlations between evolutionary rates and evolutionary time in Holometabola excluding Hymenoptera clade.** Data were retrieved from Misof *et al.* (2014). A Bayesian mixed model was performed using MCMCglmm for phylogenetic correction. *P* values were adjusted using the Benjamini & Hochberg method for multiple correction. Significant *P* values (< 0.05) for positive correlations with evolutionary time are indicated by *. BTime: branch time. Note that although *P* for nCC/nALL* is significant, but it shows negative correlation with evolutionary time.

| | Int.+Ter. | | Terminal | | Internal | |
|---|---|---|---|---|---|---|
| | ***p*** | **adjusted *p*** | ***p*** | **adjusted *p*** | ***p*** | **adjusted *p*** |
| nOP/nALL* | *0.0001 | *0.00045 | 0.43 | 0.98 | *0.0002 | *0.0018 |
| nMRP/nALL* | *0.0001 | *0.00045 | 0.28 | 0.98 | *0.037 | 0.17 |
| nCC/nALL* | 0.04 | 0.12 | 0.39 | 0.98 | 0.52 | 0.99 |
| nCRP/nALL* | 0.43 | 0.96 | 0.82 | 0.98 | 0.99 | 0.99 |
| nALL*/BTime | 0.98 | 0.99 | 0.95 | 0.98 | 0.98 | 0.99 |
| nOP/BTime | 0.92 | 0.99 | 0.95 | 0.98 | 0.97 | 0.99 |
| nMRP/BTime | 0.91 | 0.99 | 0.95 | 0.98 | 0.99 | 0.99 |
| nCC/BTime | 0.97 | 0.99 | 0.96 | 0.98 | 0.94 | 0.99 |
| nCRP/BTime | 0.99 | 0.99 | 0.98 | 0.98 | 0.99 | 0.99 |

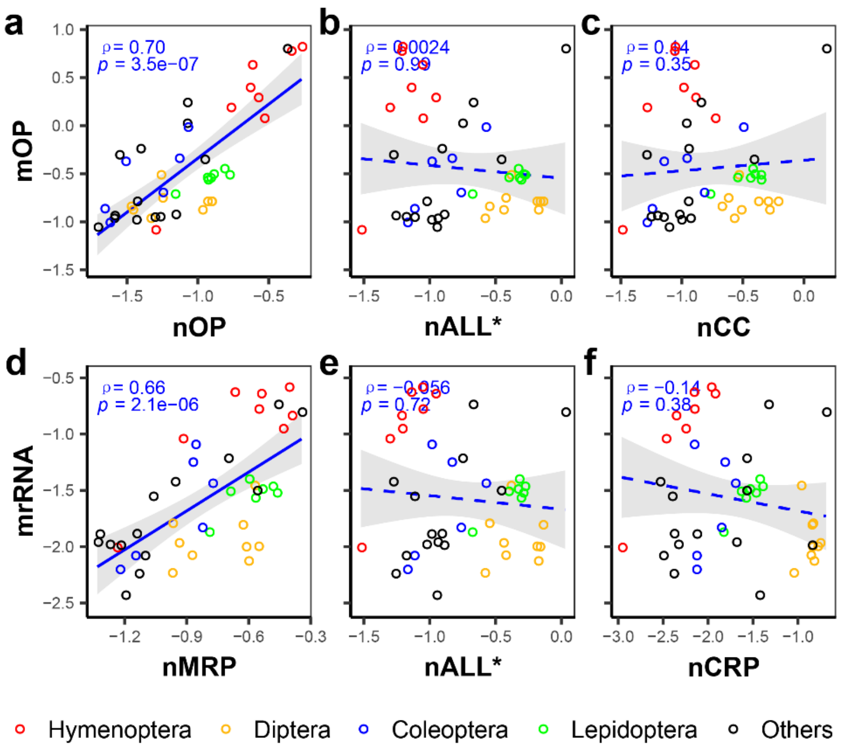


**Figure S1. Correlations between distances to HCA of mitochondrial products and nuclear encoded protein categories.** Shown are correlations of distances to the HCA for **(a)** mOP-nOP, **(b)** mOP-nALL*, **(c)** mOP-nCC, **(d)** mrRNA-nMRP, **(e)** mrRNA-nALL* and **(f)** mrRNA-nCRP. Distances to the HCA were log2 transformed. Spearman's $\rho$ and $p$ values are presented, a solid regression line indicates a significant Spearman correlation ($p < 0.05$), and dashed regression lines a non-significant correlation. Gray shadows represent 95% confidence intervals for regression lines.

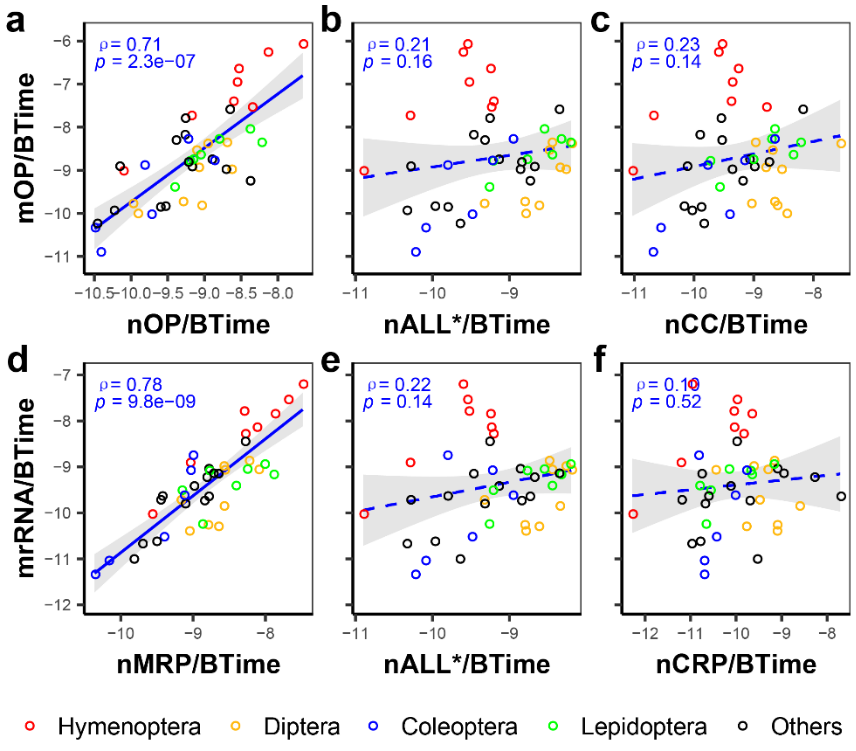


**Figure S2. Correlations between branch time (BTime) rates of mitochondrial products and nuclear encoded protein categories.** Shown are branch time rate correlations for **(a)** mOP-nOP, **(b)** mOP-nALL*, **(c)** mOP-nCC, **(d)** mrRNA-nMRP, **(e)** mrRNA-nALL* and **(f)** mrRNA-nCRP. Branch time rates were log2 transformed and Spearman's $\rho$ and $p$ values are presented. A solid regression line indicates a significant Spearman correlation ($p < 0.05$), and dashed regression line a non-significant correlation. Gray shadows represent 95% confidence intervals for regression lines.

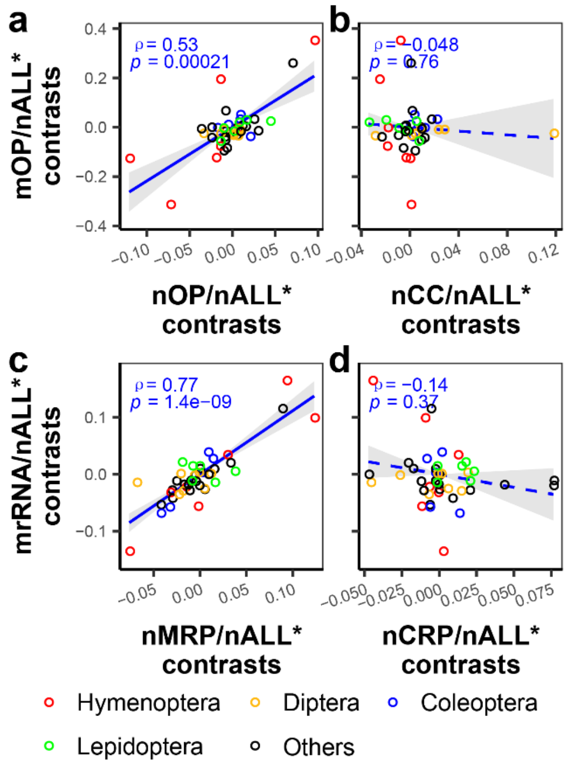


**Figure S3. Correlations between standardized contrasts of relative evolutionary rates of mitochondrial products and nuclear encoded protein categories.** Phylogenetic independent contrasts were conducted for phylogenetic correction. Shown are correlations of standardized contrasts between **(a)** mOP-nOP, **(b)** mOP-nCC, **(c)** mrRNA-nMRP and **(d)** mrRNA-nCRP. Spearman's $\rho$ and $p$ values are presented, and a solid regression line indicates a significant Spearman correlation ($p < 0.05$), and dashed regression line a non-significant correlation. Gray shadows represent 95% confidence intervals for regression lines.

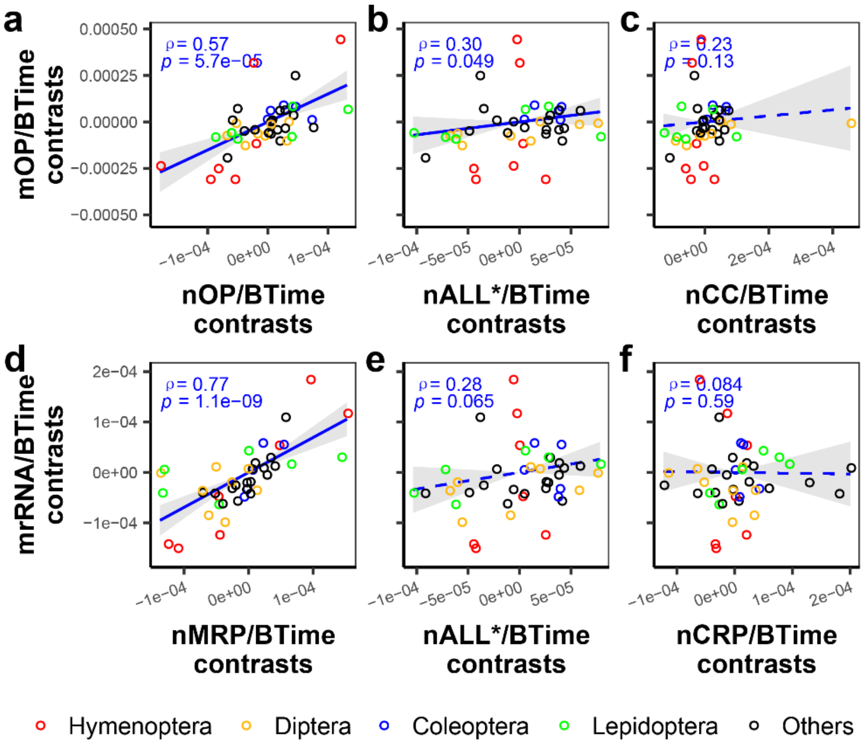


**Figure S4. Correlations between standardized contrasts of branch time evolutionary rates of mitochondrial products and nuclear encoded protein categories.** Phylogenetic independent contrasts were conducted for phylogenetic correction. Shown are correlations of standardized contrasts between **(a)** mOP-nOP, **(b)** mOP-nALL*, **(c)** mOP-nCC, **(d)** mrRNA-nMRP, (e) mrRNA-nALL* and **(f)** mrRNA-nCRP. Spearman's $\rho$ and $p$ values are presented, and a solid regression line indicates a significant Spearman correlation ($p < 0.05$), and dashed regression line a non-significant correlation. Gray shadows represent 95% confidence intervals for regression lines.

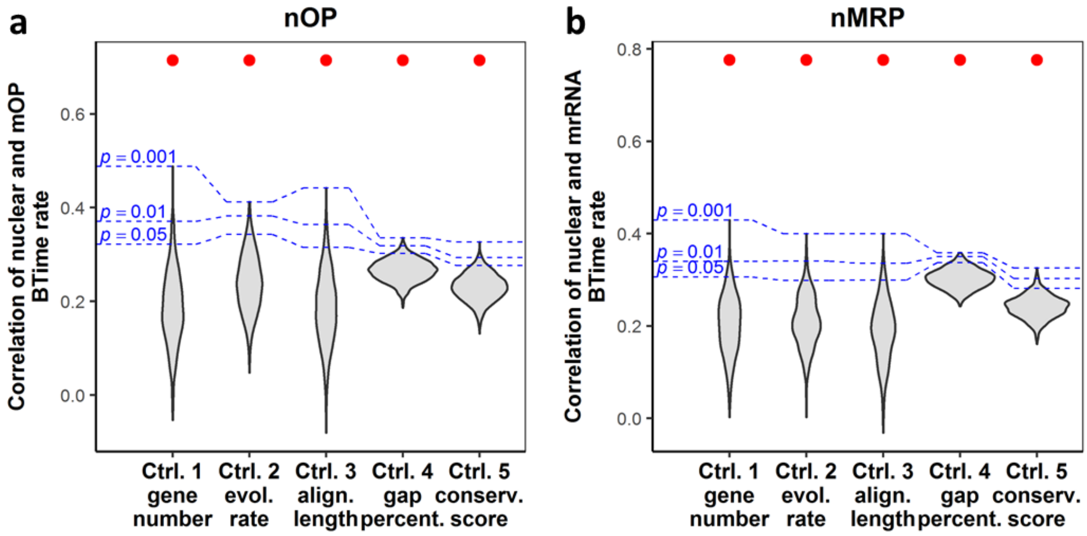


**Figure S5. Subsampling statistics for (a) nOP-mOP branch time (BTime) evolutionary rate correlation (ERC) and (b) nMRP-mrRNA ERC.** Subsampling was conducted by randomly sampling from nALL* proteins. Five different methods were used to control for gene number, evolutionary rate, alignment length, gap percentage or conservation score. As can be seen, ERCs of nOP-mOP and nMRP-mrRNA are significantly higher than ERCs for mitochondrial products and randomly generated subsamples. Red dots indicate the observed ERC of nOP-mOP or nMRP-mrRNA.

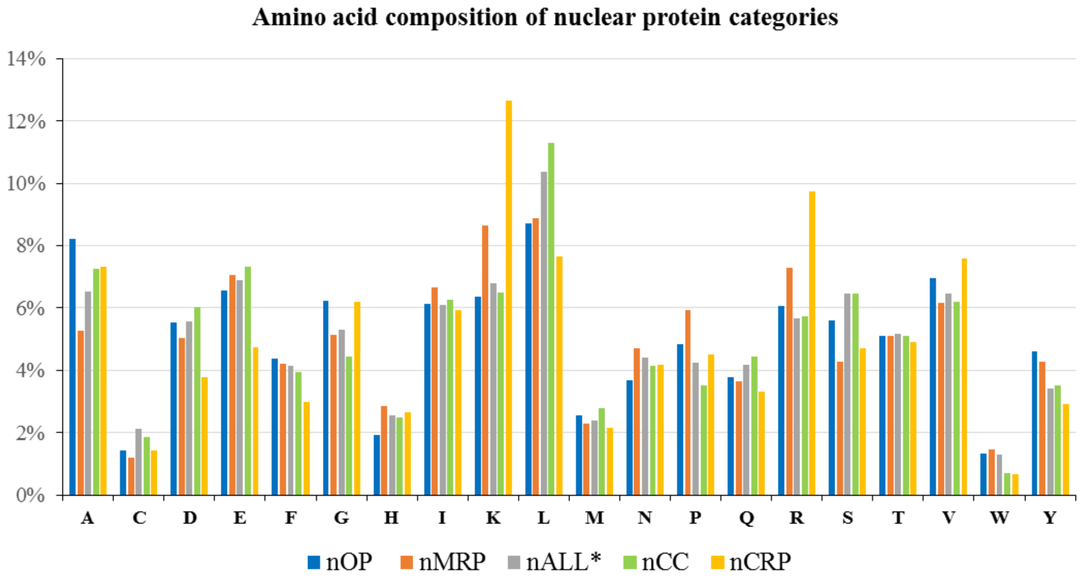


**Figure S6. Amino acid composition of different concatenated nuclear-encoded protein categories.** Each single letter presents an amino acid abbreviation. The frequencies of amino acids for nOP and nMRP do not show exceptional skews relative to other proteins.

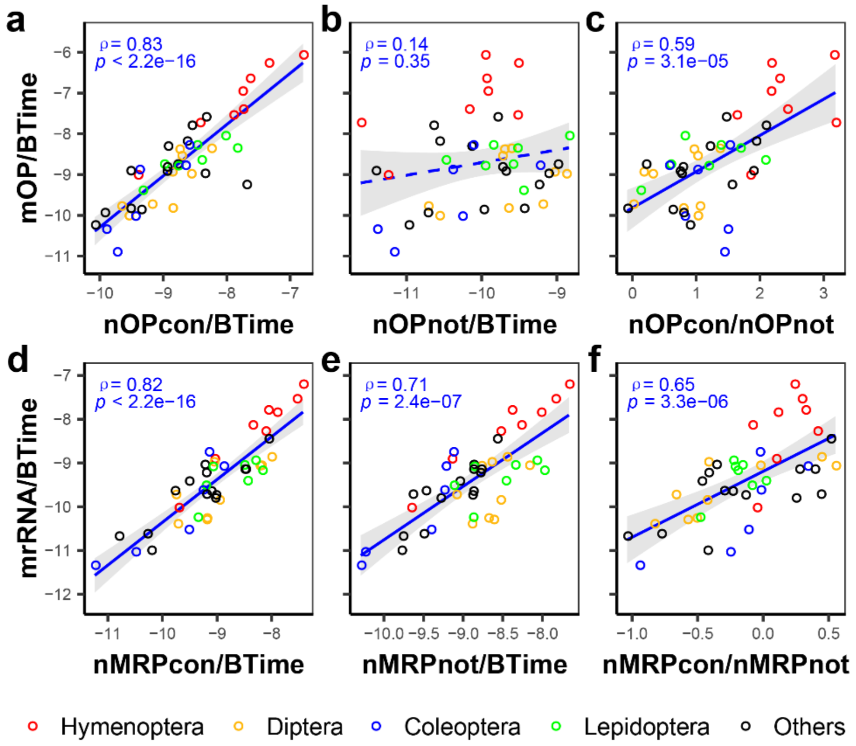


**Figure S7. Mitochondria-associated nuclear-encoded proteins in contact with mitochondrial products show stronger branch time (BTime) evolutionary rate correlations (ERCs) with mitochondrial products.** Shown are correlations between branch time rates of mOP and **(a)** nOP proteins in contact with mOP (nOPcon), **(b)** nOP proteins not in contact with mOP (nOPnot) and **(c)** ratio of nOPcon/nOPnot rates. Also shown are correlations between branch time rates of mrRNA and **(d)** nMRP amino acids in contact with mitochondrial rRNA/tRNA (nMRPcon), **(e)** nMRP amino acids not in contact with mitochondrial rRNA/tRNA (nMRPnot) and **(f)** ratio of nMRPcon/nMRPnot rates. Branch time rates and ratios were log2 transformed. Spearman's $\rho$ and $p$ values are presented, a solid regression line indicates a significant Spearman correlation ($p < 0.05$), and dashed regression line a non-significant correlation. Gray shadows represent 95% confidence intervals for regression lines.

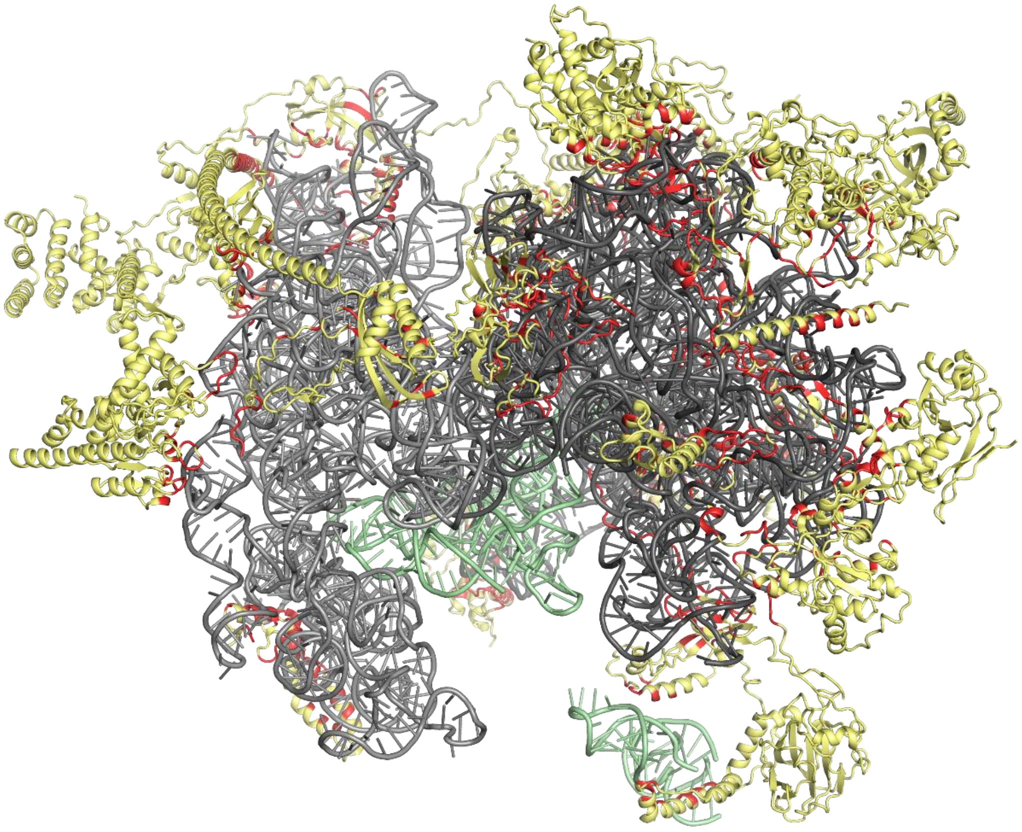

**Figure S8. Structure of mammalian mitochondrial ribosome.** Twenty-five nMRP proteins in Misof *et al.* (2014) data which have mammalian orthologs, are presented in yellow. Mitochondrial rRNAs are colored gray. Mitochondrial tRNAs are colored green. Amino acids in contact with mitochondrial rRNAs or tRNAs are colored red. Structure information was retrieved from Greber *et al.* (2015) and redrawn.

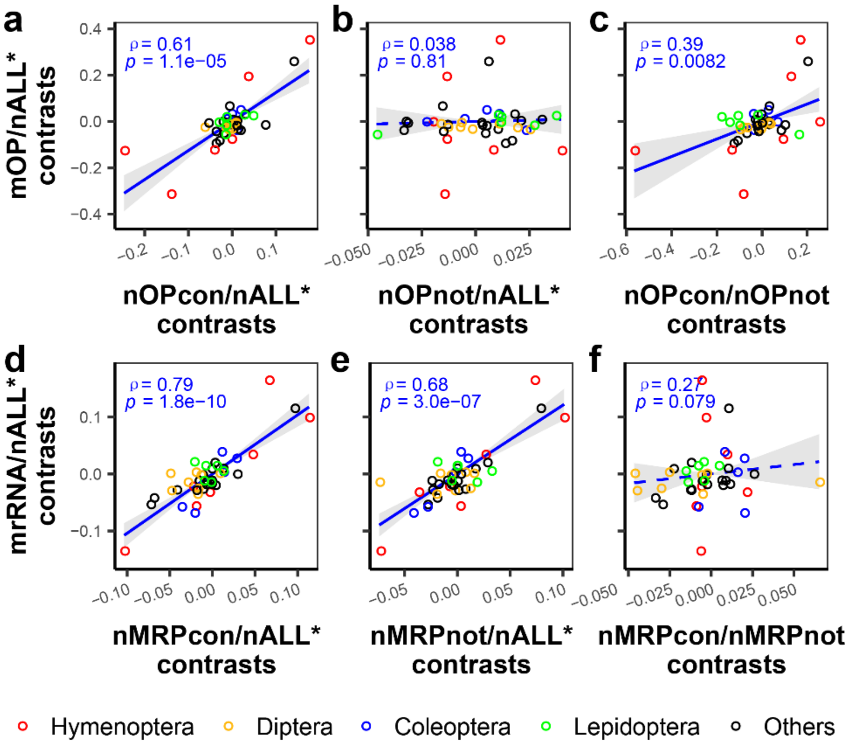


**Figure S9. Correlations between standardized contrasts of relative evolutionary rates of mitochondrial products and nuclear encoded protein categories in contact or not in contact with mitochondrial products.** Phylogenetic independent contrasts were conducted for phylogenetic correction. Shown are standardized contrast correlations between relative rates of mOP and **(a)** nOPcon, **(b)** nOPnot and **(c)** ratio nOPcon/nOPnot. Also shown are standardized contrast correlations between relative rates of mrRNA and **(d)** nMRPcon, **(e)** nMRPnot and **(f)** ratio nMRPcon/nMRPnot. Spearman's $\rho$ and $p$ values are presented, solid regression line indicates a significant Spearman correlation ($p < 0.05$), and dashed regression line a non-significant correlation. Gray shadows represent 95% confidence intervals for regression lines.

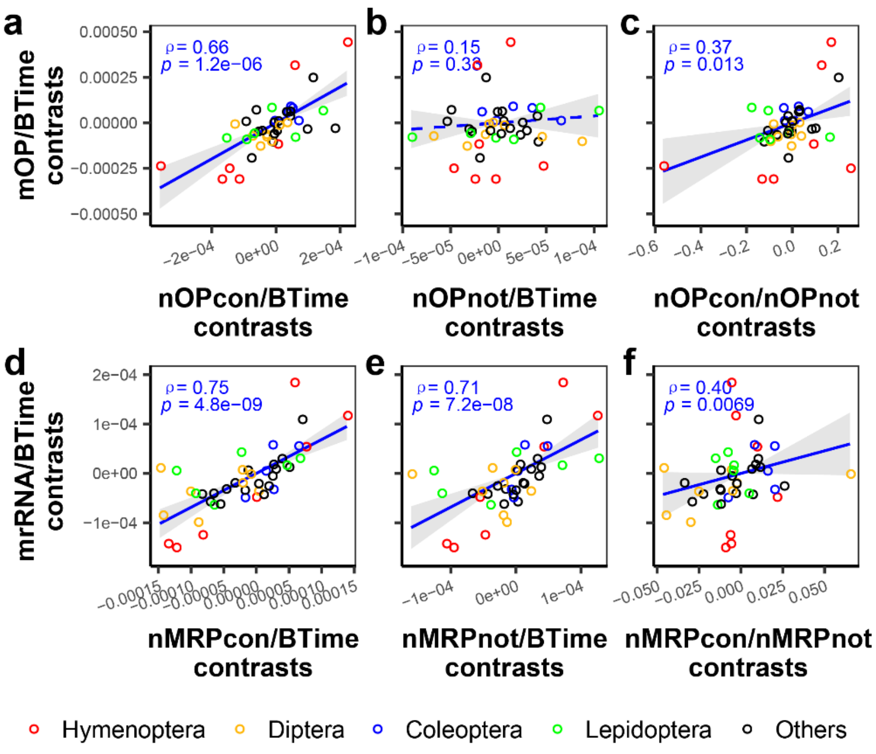


**Figure S10. Correlations between standardized contrasts of branch time rates of mitochondrial products and nuclear encoded protein categories in contact or not in contact with mitochondrial products.** Phylogenetic independent contrasts were conducted for phylogenetic correction. Shown are standardized contrast correlations between branch time rates of mOP and **(a)** nOPcon, **(b)** nOPnot and **(c)** ratio nOPcon/nOPnot. Also shown are standardized contrast correlation between branch time rates of mrRNA and **(d)** nMRPcon, **(e)** nMRPnot and **(f)** ratio nMRPcon/nMRPnot. Spearman's $\rho$ and $p$ values are presented, a solid regression line indicates a significant Spearman correlation ($p < 0.05$), and dashed regression line a non-significant correlation. Gray shadows represent 95% confidence intervals for regression lines.

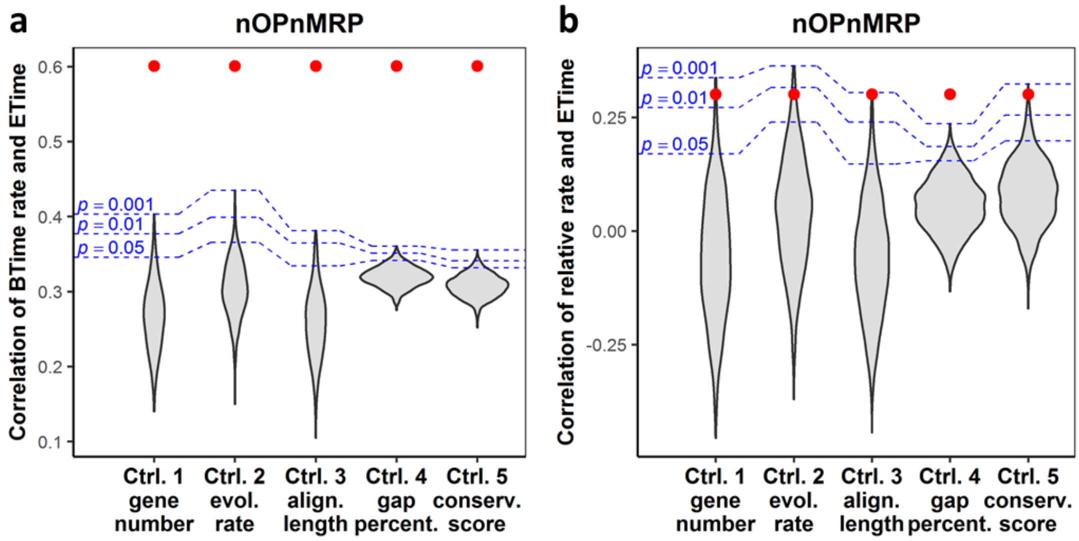


**Figure S11. Subsampling statistics for correlations between evolutionary rates of nOPnMRP and their evolutionary time.** Subsampling statistics are shown for **(a)** correlations between branch time rate of nOPnMRP and their evolutionary time, and **(b)** correlations between relative rate of nOPnMRP and their evolutionary time. Estimated dates at the midpoints of each branch were used as evolutionary time. Evolutionary rates on both terminal and internal branches are included. Subsampling was conducted by randomly sampling from nALL* proteins. Five different analyses were used to control for gene number, evolutionary rate, alignment length, gap percentage or conservation score. Red dots indicate the observed correlations between evolutionary rates of nOPnMRP and evolutionary time. As can be seen, evolutionary rates of nOPnMRP show significantly higher correlations with evolutionary time than those of randomly generated samples.

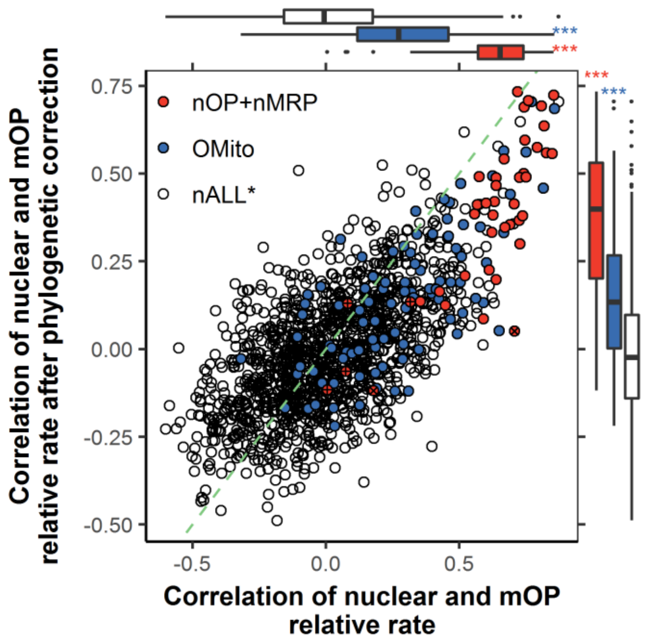


**Figure S12. Correlations of nuclear and mOP relative rates before and after phylogenetic correction.** Red indicates nOP and nMRP proteins, + indicates 4 nOP proteins not in contact with mOP, × indicates two nOP proteins whose mOP contacting information is unknown. Blue indicates other mitochondria-associated proteins identified by GO/MitoDrome annotations (OMito). Green dashed line represents the diagonal line. Asterisks in marginal boxplots indicate significant difference from nALL*; ns indicates no significant difference ($p > 0.05$); * $p < 0.05$, ** $p < 0.01$; *** $p < 0.001$. As can been seen, phylogenetic correction reduces the significance and changes ranks of correlations.

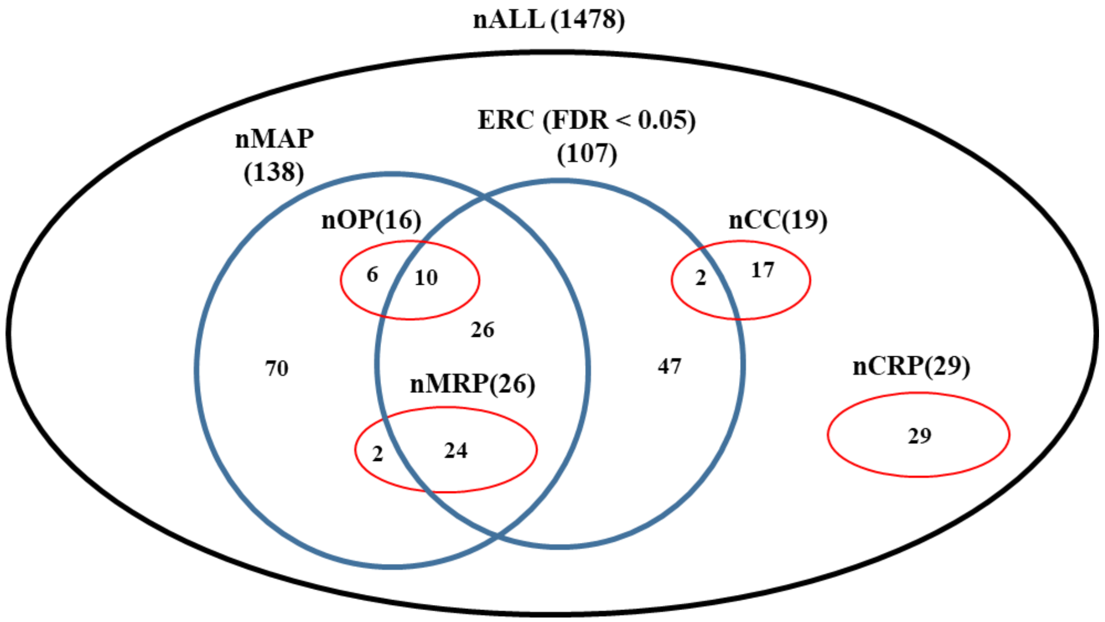


**Figure S13. Venn graph of different nuclear-encoded protein categories.** nALL: all 1478 nuclear-encoded single copy genes from Misof *et al.* (2014). nMAP: 138 mitochondria-associated proteins identified by the GOstats or in MitoDrome database. nALL: 1478 single copy nuclear-encoded proteins from Misof *et al.* (2014). ERC (FDR < 0.05) refers to 107 single copy nuclear-encoded proteins with significant evolutionary rate correlations to mOP (FDR < 0.05). For other abbreviations see Table 1.

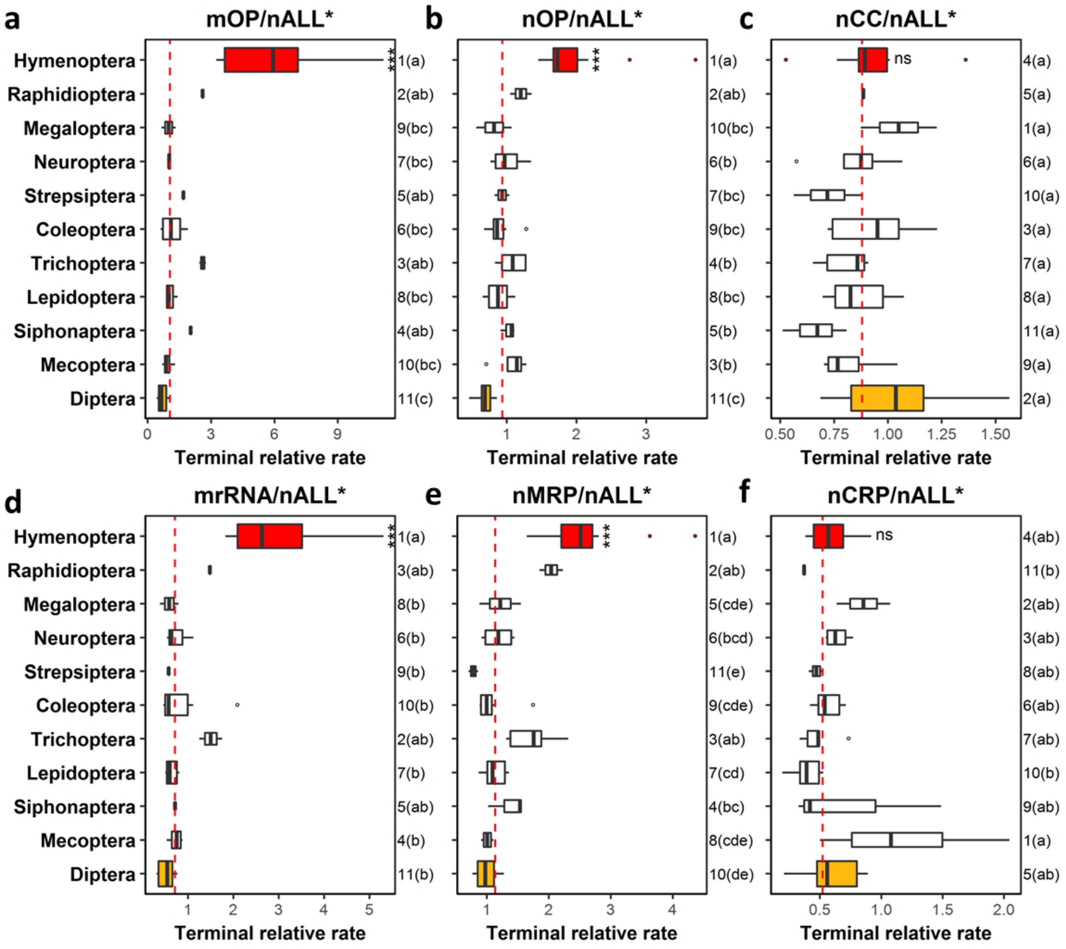


**Figure S14. Relative evolutionary rates of concatenated mitochondrial products and nuclear protein categories vary across taxa in Holometabola**. Relative evolutionary rates of **(a)** mitochondrial-encoded OXPHOS proteins (mOP), **(b)** nuclear-encoded OXPHOs proteins (nOP), **(c)** nuclear-encoded cell cycle proteins (nCC), **(d)** mitochondrial ribosomal RNAs (mrRNA), **(e)** nuclear-encoded mitochondrial ribosomal proteins (nMRP), and **(f)** nuclear-encoded cytosolic ribosomal proteins (nCRP). Numbers on right indicate ranks of each order. Orders sharing the same letter(s) in parentheses have no significant difference from each other ($p < 0.05$). Asterisks on Hymenoptera indicate significant difference from Diptera using Wilcoxon rank sum test; ns indicates no significant difference ($p > 0.05$); * $p < 0.05$; ** $p < 0.01$; *** $p < 0.001$.

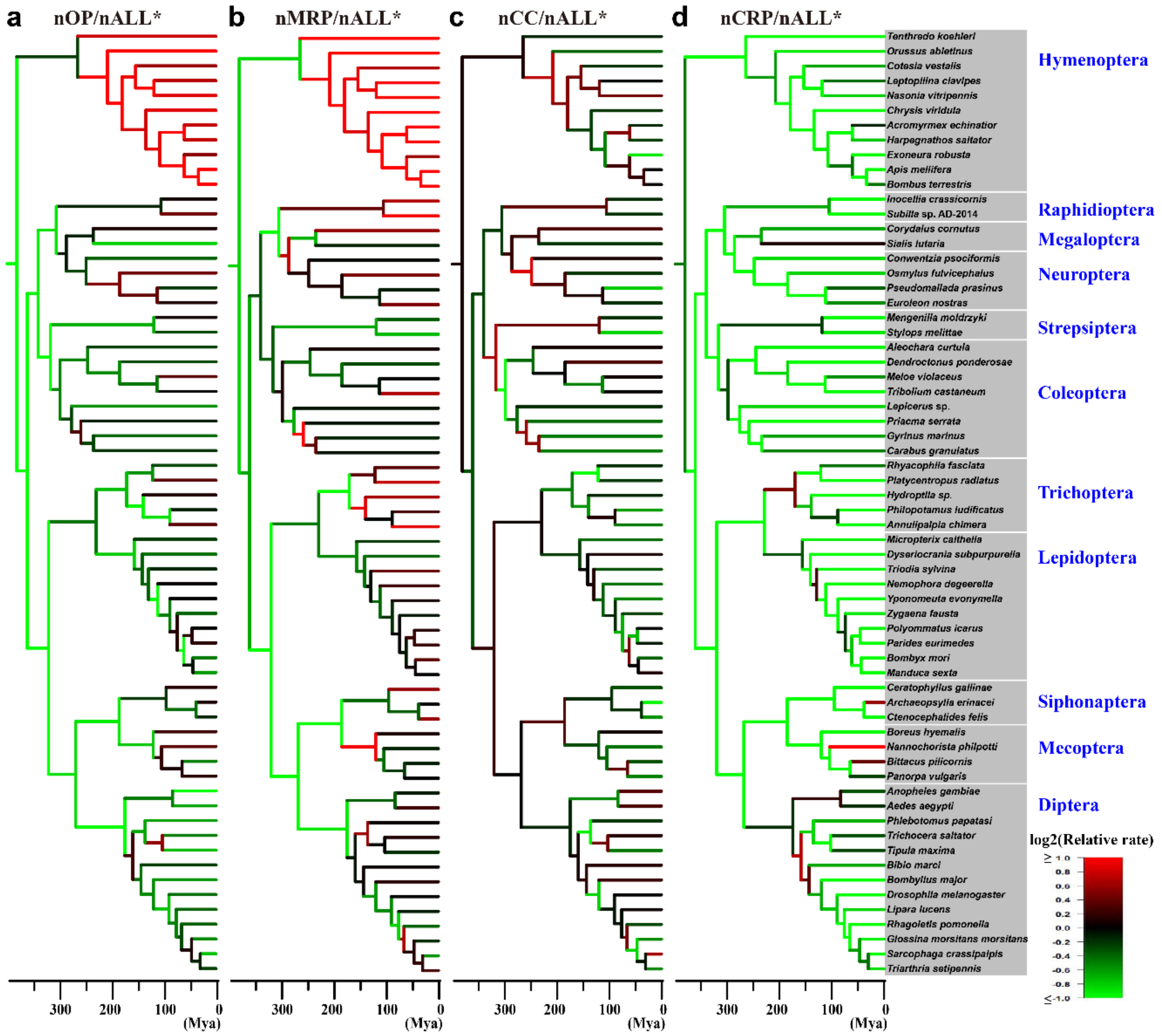


**Figure S15. Dated phylogenetic trees in Holometabola.** These dated trees were retrieved from Misof *et al.* (2014). Branches were colored based on their relative evolutionary rates of **(a)** nOP, **(b)** nMRP, **(c)** nCC and **(d)** nCRP. Relative evolutionary rates were log2 transformed, then mapped to color gradient from green (low) to red (high).

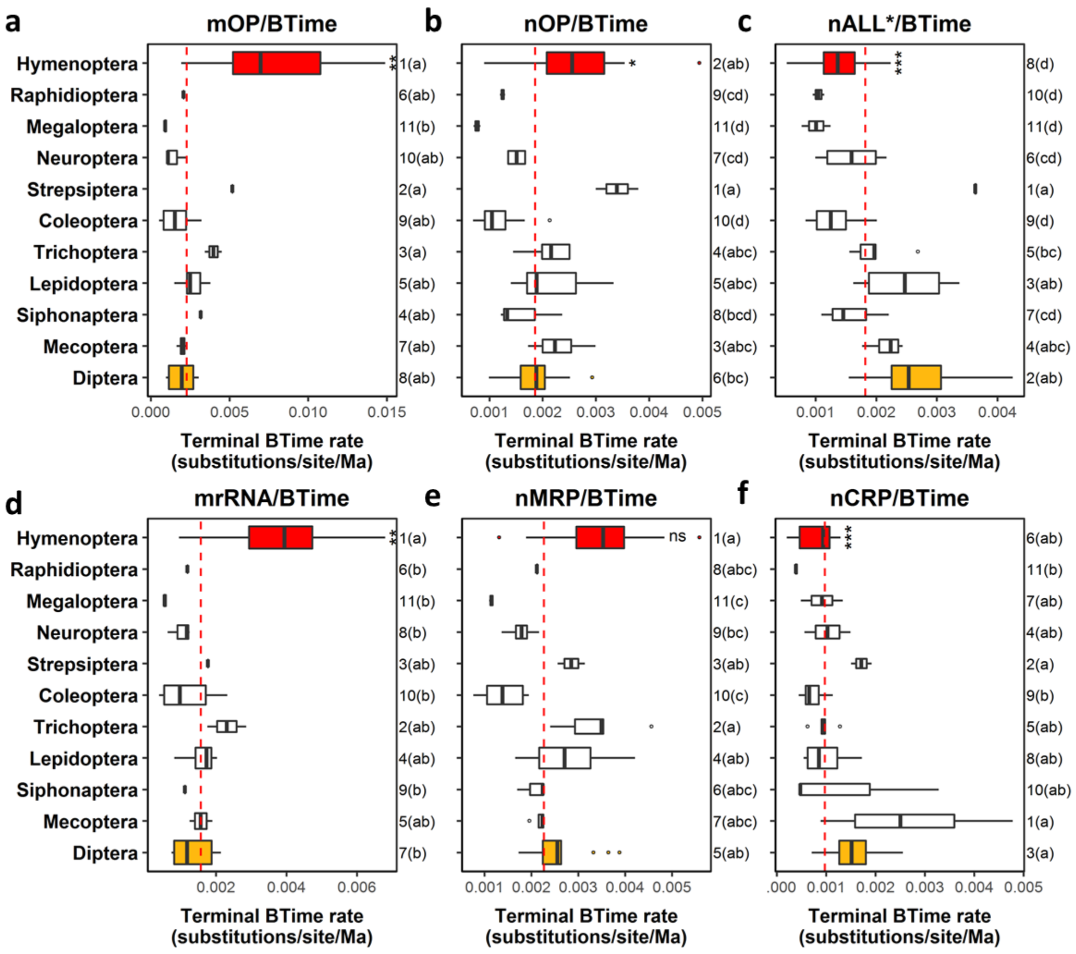


**Figure S16. Branch time (BTime) evolutionary rates of concatenated mitochondrial products and nuclear protein categories vary across insect orders in Holometabola**. Branch time evolutionary rates of **(a)** mitochondrial-encoded OXPHOS proteins (mOP), **(b)** nuclear-encoded OXPHOs proteins (nOP), **(c)** all 1478 nuclear-encoded proteins excluding these with mitochondrial GO annotations (nALL*), **(d)** mitochondrial ribosomal RNAs (mrRNA), **(e)** nuclear encoded mitochondrial ribosomal proteins (nMRP), and **(f)** nuclear encoded cytosolic ribosomal proteins (nCRP). Numbers on right indicate ranks of each order. Orders sharing the same letter(s) in parentheses have no significant difference from each other ($p < 0.05$). Asterisks on Hymenoptera indicate significant difference from Diptera using Wilcoxon rank sum test; ns indicates no significant difference ($p > 0.05$); * $p < 0.05$; ** $p < 0.01$; *** $p < 0.001$.

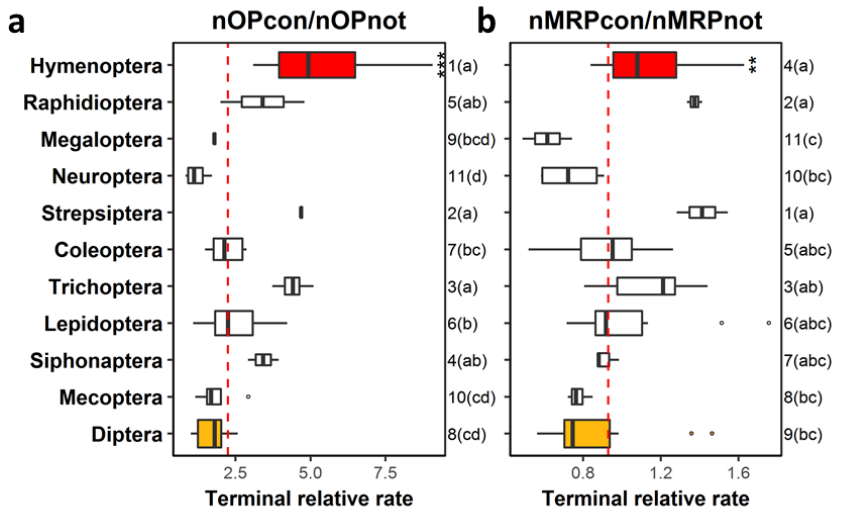


**Figure S17. Evolutionary rate ratios of components in contact over those not in contact with mitochondrial products in Holometabola. (a)** Ratios of nOP proteins in contact with mOP (nOPcon) over those not in contact (nOPnot). **(b)** Ratios of nMRP amino acids in contact with mitochondrial rRNA/tRNA (nMRPcon) over those not in contact (nMRPnot). The orders are ranked based on their median values. Orders sharing the same letter(s) have no significant difference from each other ($p < 0.05$). Asterisks on Hymenoptera indicate significant difference from Diptera using Wilcoxon rank sum test; ns indicates no significant difference ($p > 0.05$); * $p < 0.05$; ** $p < 0.01$; *** $p < 0.001$.

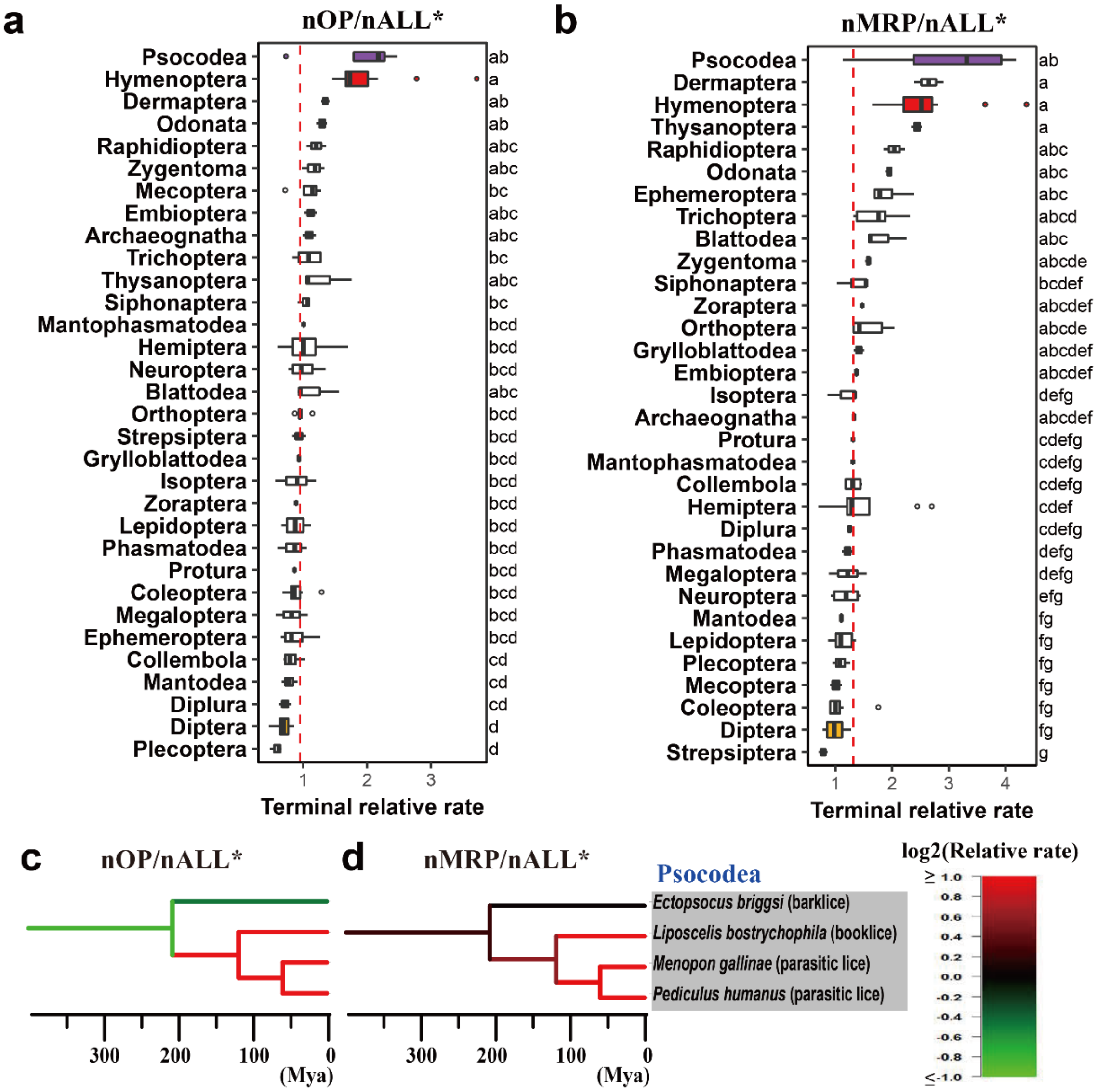


**Figure S18. Relative rates of mitochondria-associated nuclear-encoded proteins show acceleration within Psocodea.** Concatenated alignments were used for estimating evolutionary rates of terminal branches for **(a)** nOP and **(b)** nMRP in Hexapoda. Insect orders are ranked based on their median values. Orders sharing the same letter(s) have no significant difference from each other ($p < 0.05$). Also shown are dated phylogenetic trees in Psocodea with branches colored based on relative evolutionary rates of **(c)** nOP and **(d)** nMRP.

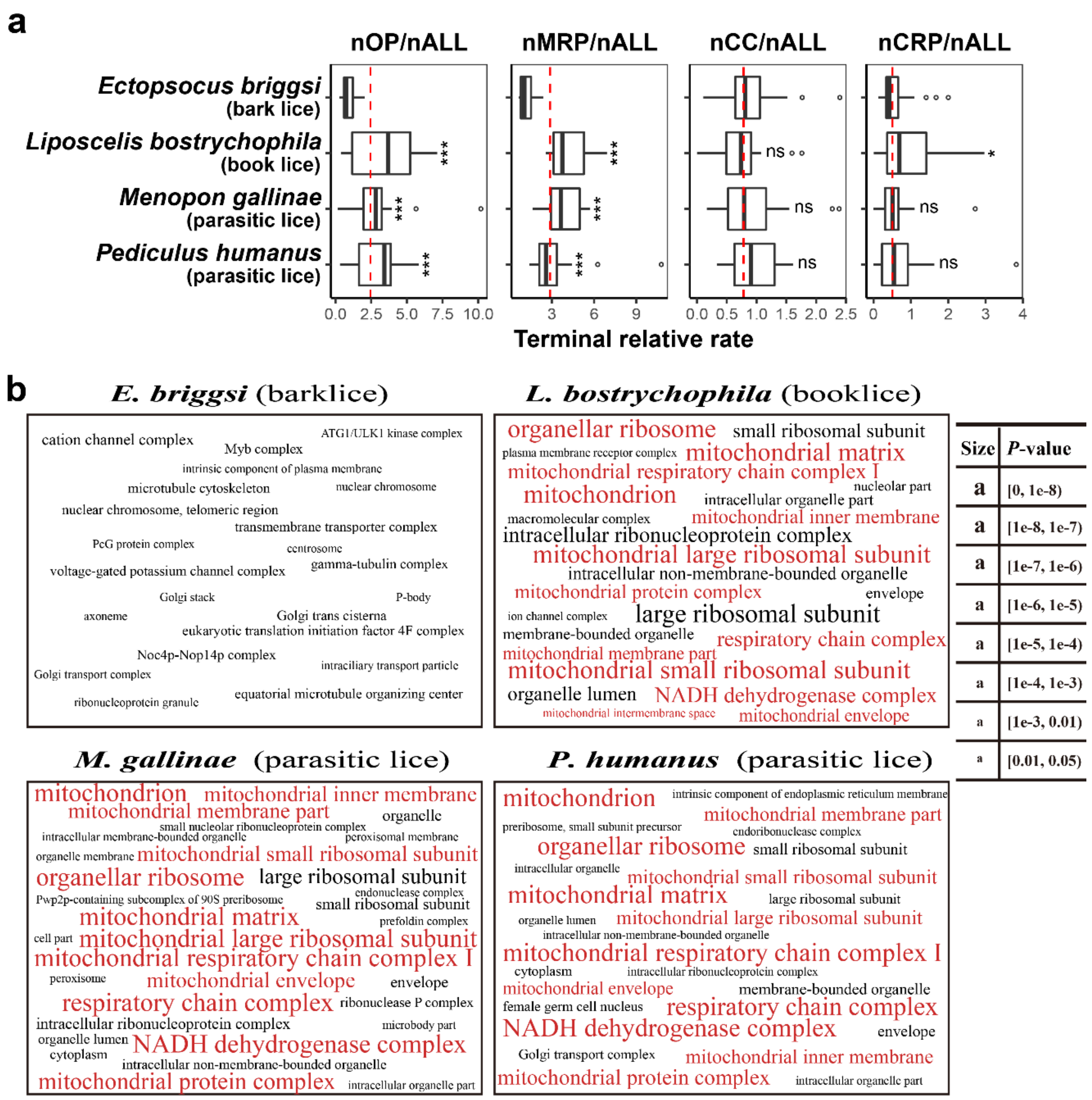


**Figure S19. Relative rate variation of mitochondria-associated nuclear-encoded proteins in Psocodea. (a)** Using individual gene analyses**,** relative evolutionary rates of nuclear gene categories within the Psocodea are compared. Asterisks indicate significant differences from *Ectopsocus briggsi*; ns indicates no significant difference ($p > 0.05$); * $p < 0.05$; ** $p < 0.01$; *** $p < 0.001$. **(b)** Word clouds of overrepresented GO items in the top 10% fastest evolving proteins from each Psocodea species. Mitochondria-related GO items are colored in red.

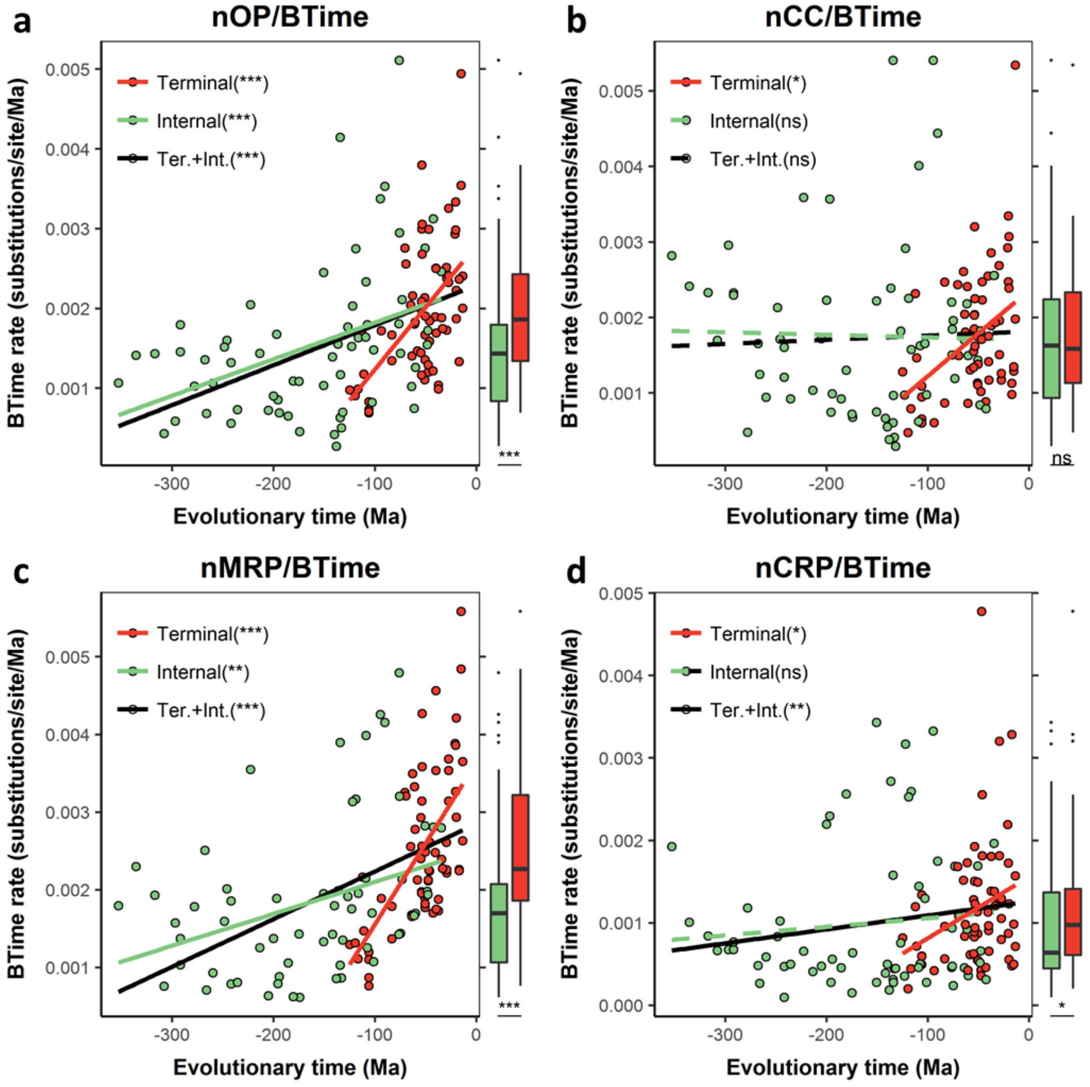


**Figure S20. Correlations between branch time (BTime) evolutionary rates and their estimated evolutionary time in Holometabola.** Shown are correlations between evolutionary time and branch time rates for **(a)** nOP, **(b)** nALL*, **(c)** nMRP and **(d)** nCRP in Holometabola. The dated phylogeny is from Misof *et al.* (2014), and estimated dates at the midpoints of each branch was chosen for analysis. Red indicates terminal branches. Green indicates internal branches. Asterisks in parenthesis indicate Spearman correlation significance; ns indicates no significant difference ($p > 0.05$); * $p < 0.05$; ** $p < 0.01$; *** $p < 0.001$.

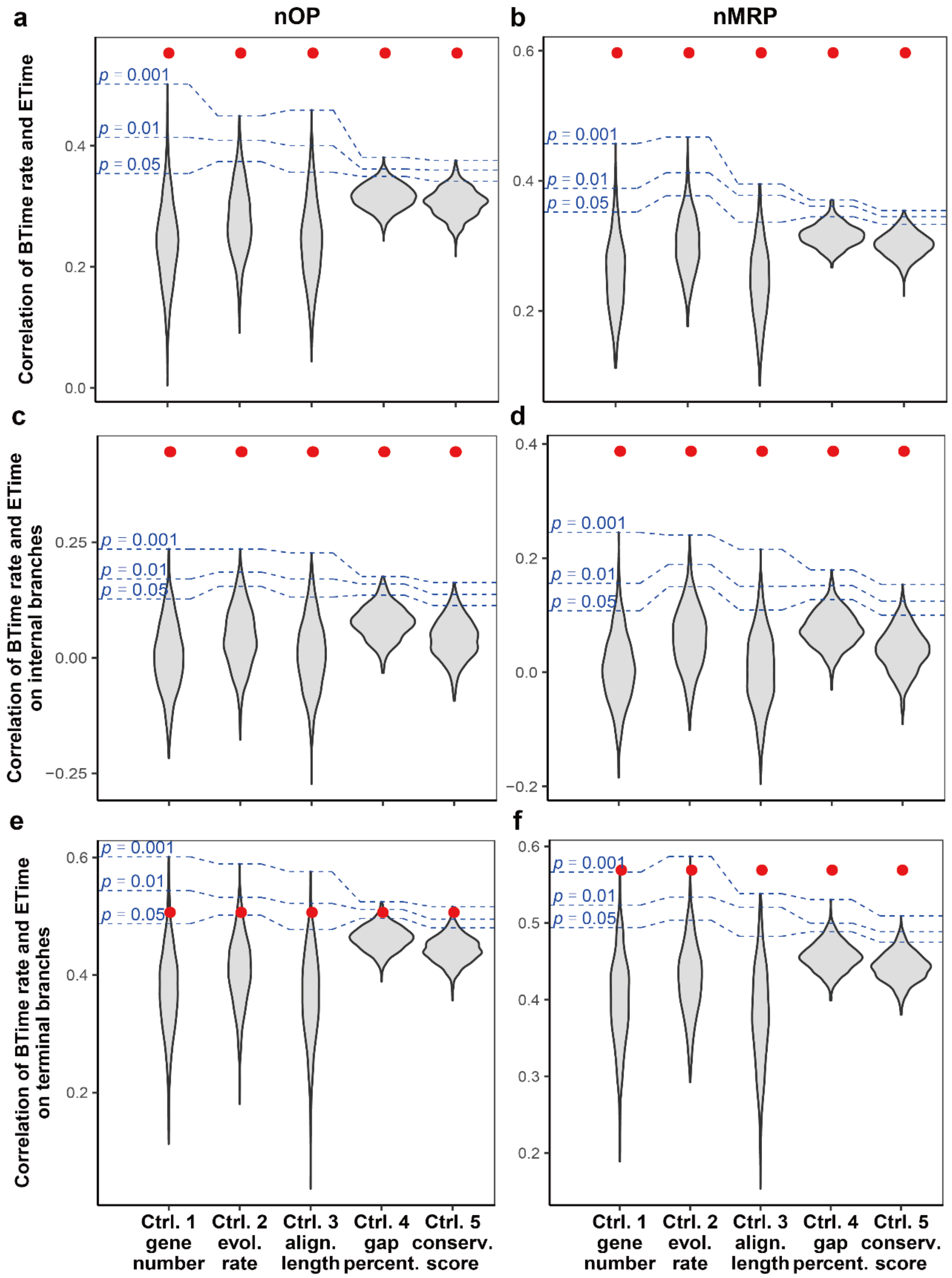


**Figure S21. Subsampling statistics for correlations between branch time (BTime) evolutionary rates and their evolutionary time (ETime).** Subsampling statistics for correlations between branch time rate of nOP and evolutionary time on **(a)** terminal plus internal branches, **(c)** internal branches or **(e)** terminal branches only. Subsampling statistics for correlations between branch time rate of nMRP and evolutionary time on **(b)** terminal plus internal branches, **(d)** internal branches or **(f)** terminal branches only. Estimated dates at the midpoints of each branch were used as evolutionary time. Subsampling was conducted by randomly sampling from nALL*

proteins. Subsampling was done to control for gene number, evolutionary rate, alignment length, gap percentage or conservation score. Red dots indicate observed correlations between branch time rates of nOP or nMRP and evolutionary time. As can be seen, evolutionary rates of nOP and nMRP both show significantly higher correlations with evolutionary time than those of randomly generated subsamples.

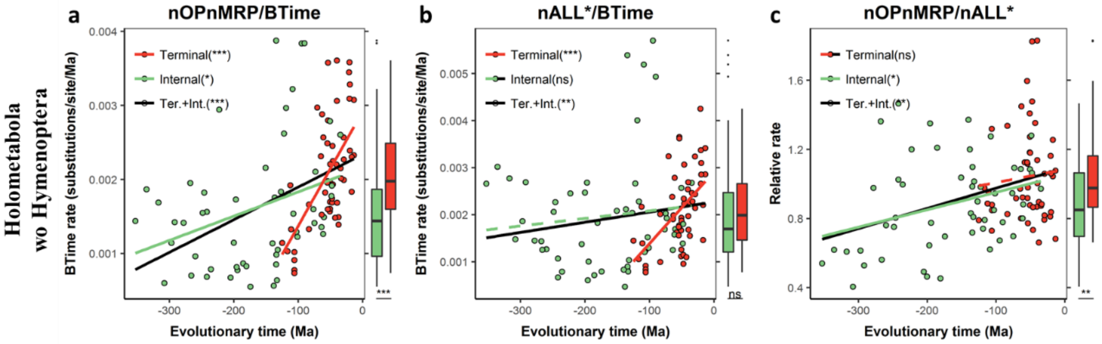


**Figure S22. Correlations between evolutionary rates and their estimated evolutionary time in Holometabola excluding Hymenoptera clade.** Shown are correlations between evolutionary time and branch time rates of **(a)** nOPnMRP, **(b)** nALL* and **(c)** relative rate of nOPnMRP in Holometabola excluding the Hymenoptera clade. Branch dates are from Misof *et al.* (2014), with each branch midpoint used for the analysis. Red indicates terminal branches. Green indicates internal branches. Asterisks in parenthesis indicate Spearman correlation significance; ns indicates not significant ($p > 0.05$); * $p < 0.05$; ** $p < 0.01$; *** $p < 0.001$.

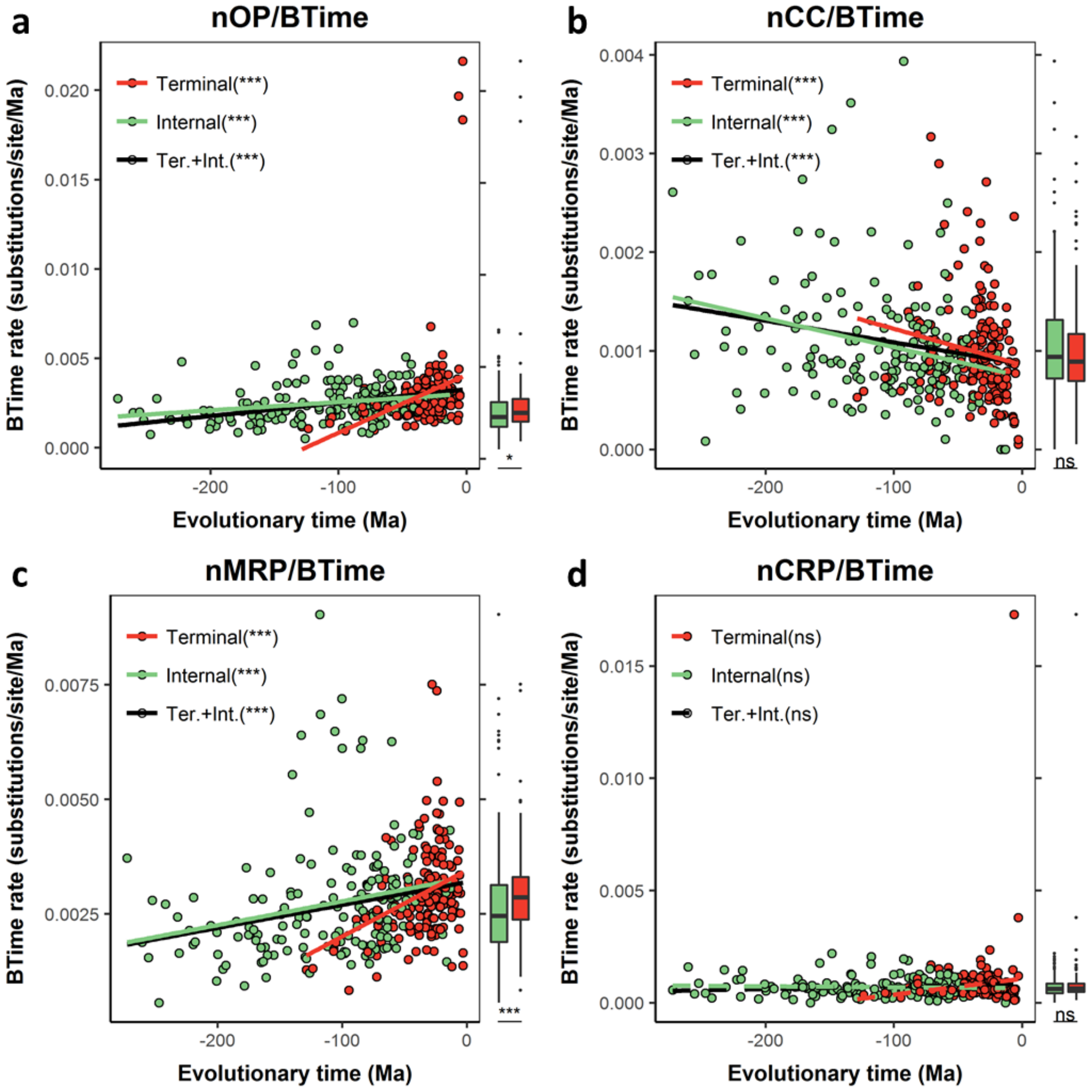


**Figure S23. Correlations between branch time (BTime) evolutionary rates and their estimated evolutionary time in Hymenoptera.** Shown are correlations between evolutionary time and branch time rates of **(a)** nOP, **(b)** nALL*, **(c)** nMRP and **(d)** nCRP in Hymenoptera. Branch dates are from Misof *et al.* (2014), with each branch midpoint used for the analysis. Red indicates terminal branches. Green indicates internal branches. Asterisks in parenthesis indicate Spearman correlation significance; ns indicates not significant ($p > 0.05$); * $p < 0.05$; ** $p < 0.01$; *** $p < 0.001$.

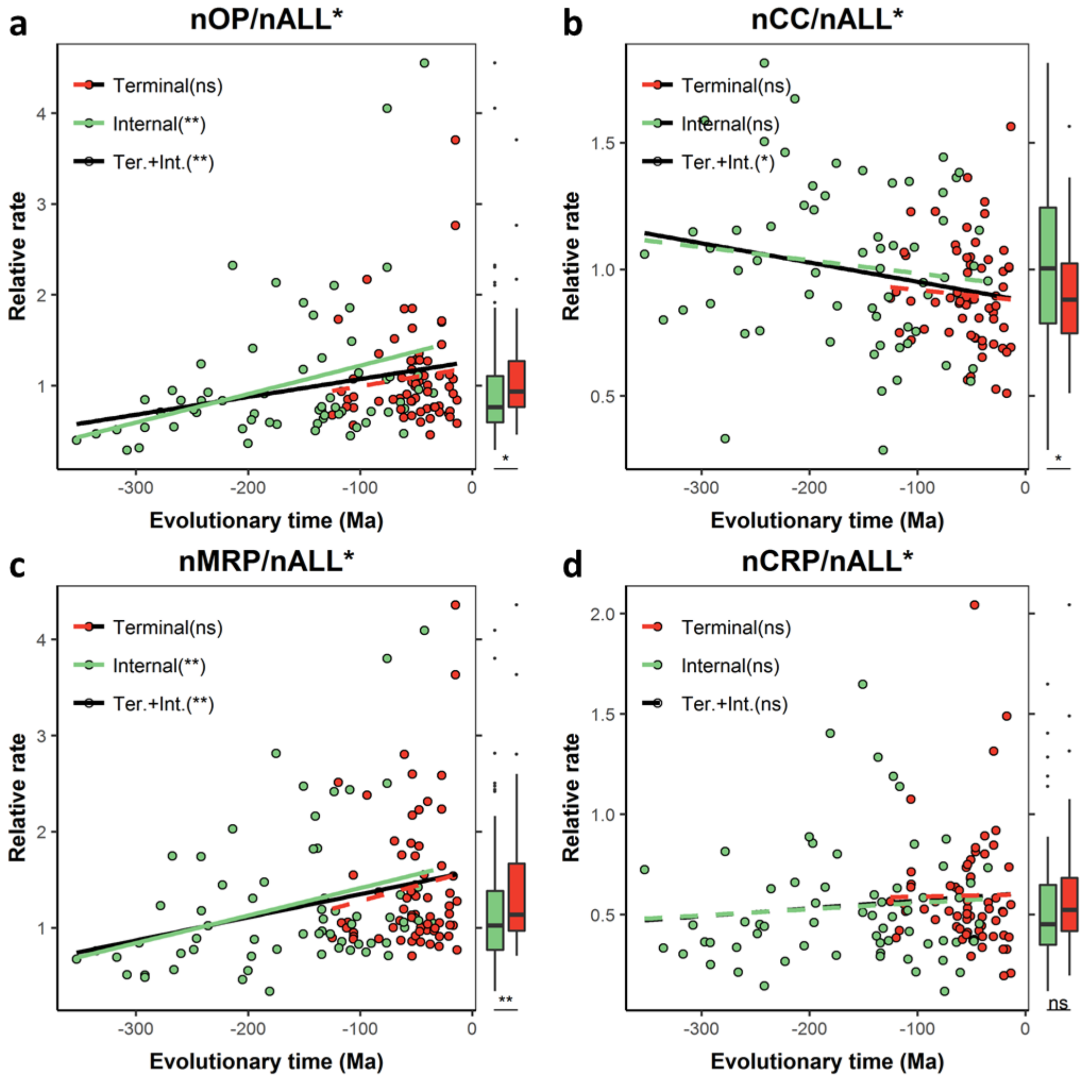


**Figure S24. Correlations between relative evolutionary rates and their estimated evolutionary time in Holometabola.** Shown are correlations between evolutionary time and relative evolutionary rates of **(a)** nOP, **(b)** nCC, **(c)** nMRP and **(d)** nCRP in Holometabola. Branch dates are from Misof *et al.* (2014), with each branch midpoint used for the analysis Red indicates terminal branches. Green indicates internal branches. Asterisks in parenthesis indicate Spearman correlation significance; ns indicates not significant (p > 0.05); * $p < 0.05$; ** $p < 0.01$; *** $p < 0.001$.

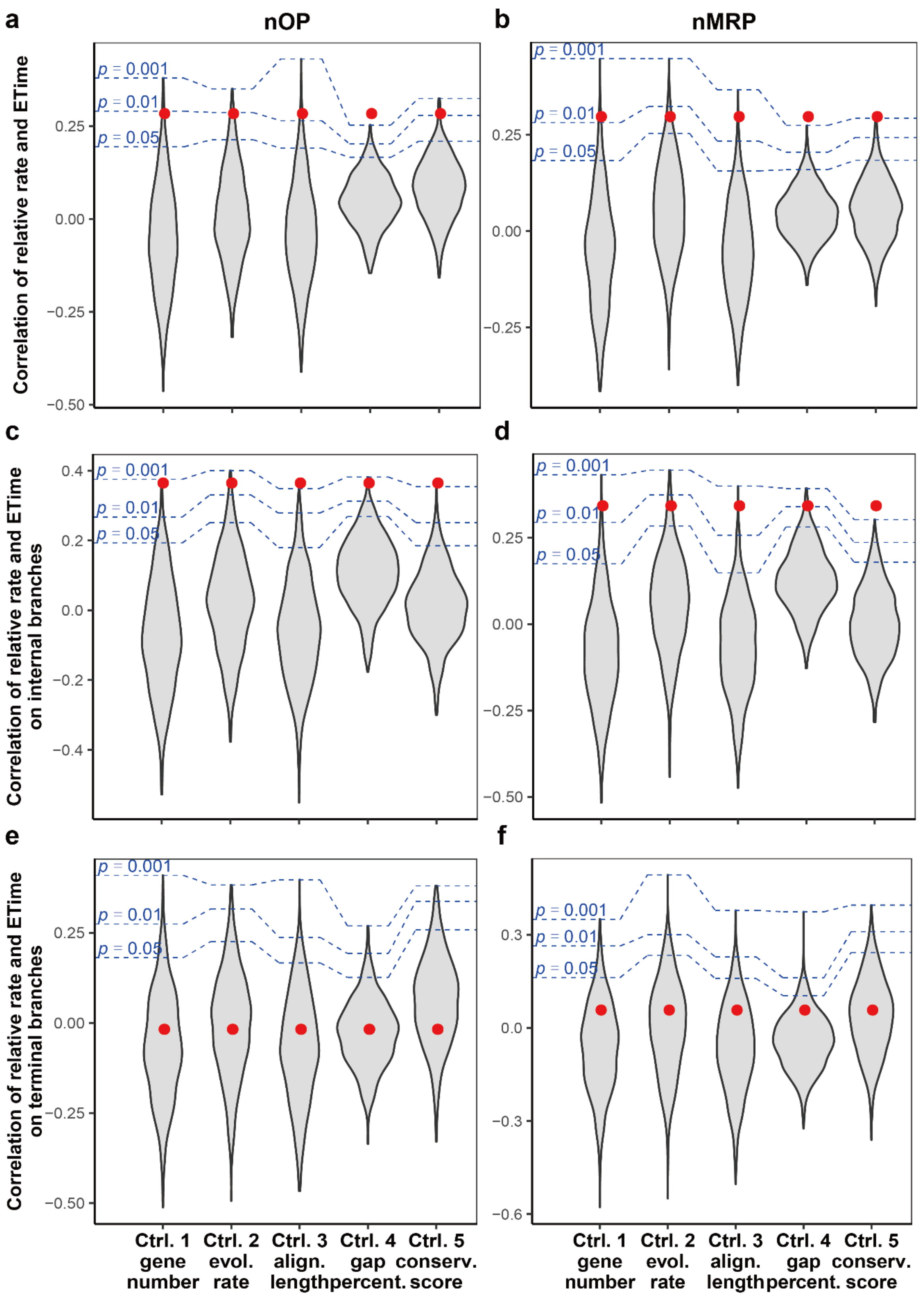


**Figure S25. Subsampling statistics for correlations between relative evolutionary rates and their evolutionary time (ETime).** Subsampling statistics for correlations between branch time rate of nOP and evolutionary time on **(a)** terminal plus internal branches, **(c)** internal branches or **(e)** terminal branches only. Subsampling statistics for correlations between branch time rate of nMRP and evolutionary time on **(b)** terminal plus internal branches, **(d)** internal branches or **(f)** terminal branches only. Estimated dates at the midpoints of each branch were used as evolutionary time. Subsampling was conducted by randomly sampling from nALL* proteins. Subsampling was done to control for gene number, evolutionary rate, alignment length, gap percentage or conservation score. Red dots indicate observed correlations between branch time rates of nOP or nMRP and evolutionary time. As can be seen, evolutionary rates of nOP and nMRP both show significantly higher correlations with evolutionary time than those of randomly generated subsamples, except for evolutionary rates on terminal branches.

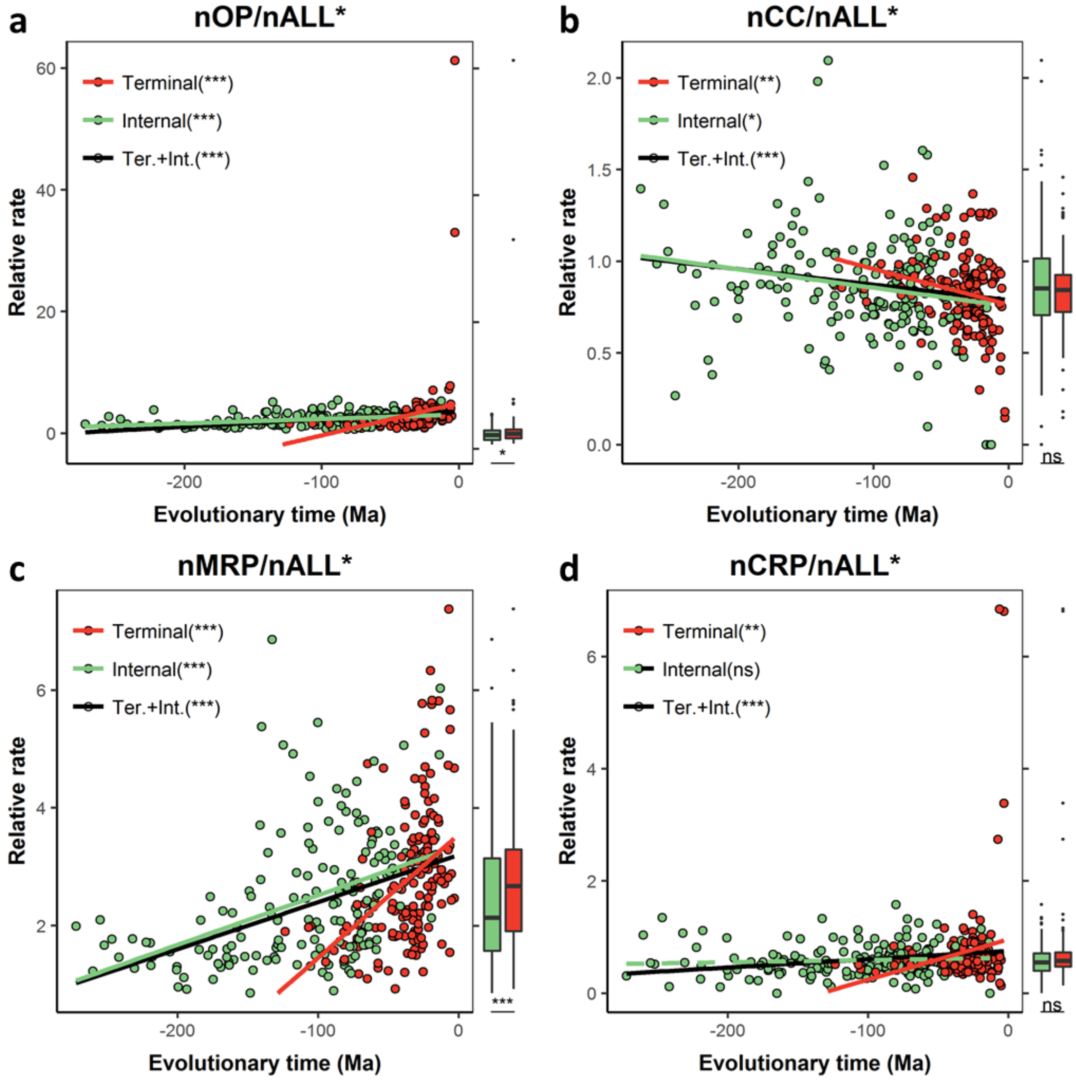


**Figure S26. Correlations between relative evolutionary rates and their estimated evolutionary time in Hymenoptera.** Shown are correlations between evolutionary time and relative evolutionary rates of **(a)** nOP, **(b)** nCC, **(c)** nMRP and (d) nCRP in Hymenoptera. Branch dates are from Misof *et al.* (2014), with each branch midpoint used for the analysis. Red indicates terminal branches. Green indicates internal branches. Asterisks in parenthesis indicate Spearman correlation significance; ns indicates not significant ($p > 0.05$); * $p < 0.05$; ** $p < 0.01$; *** $p < 0.001$.

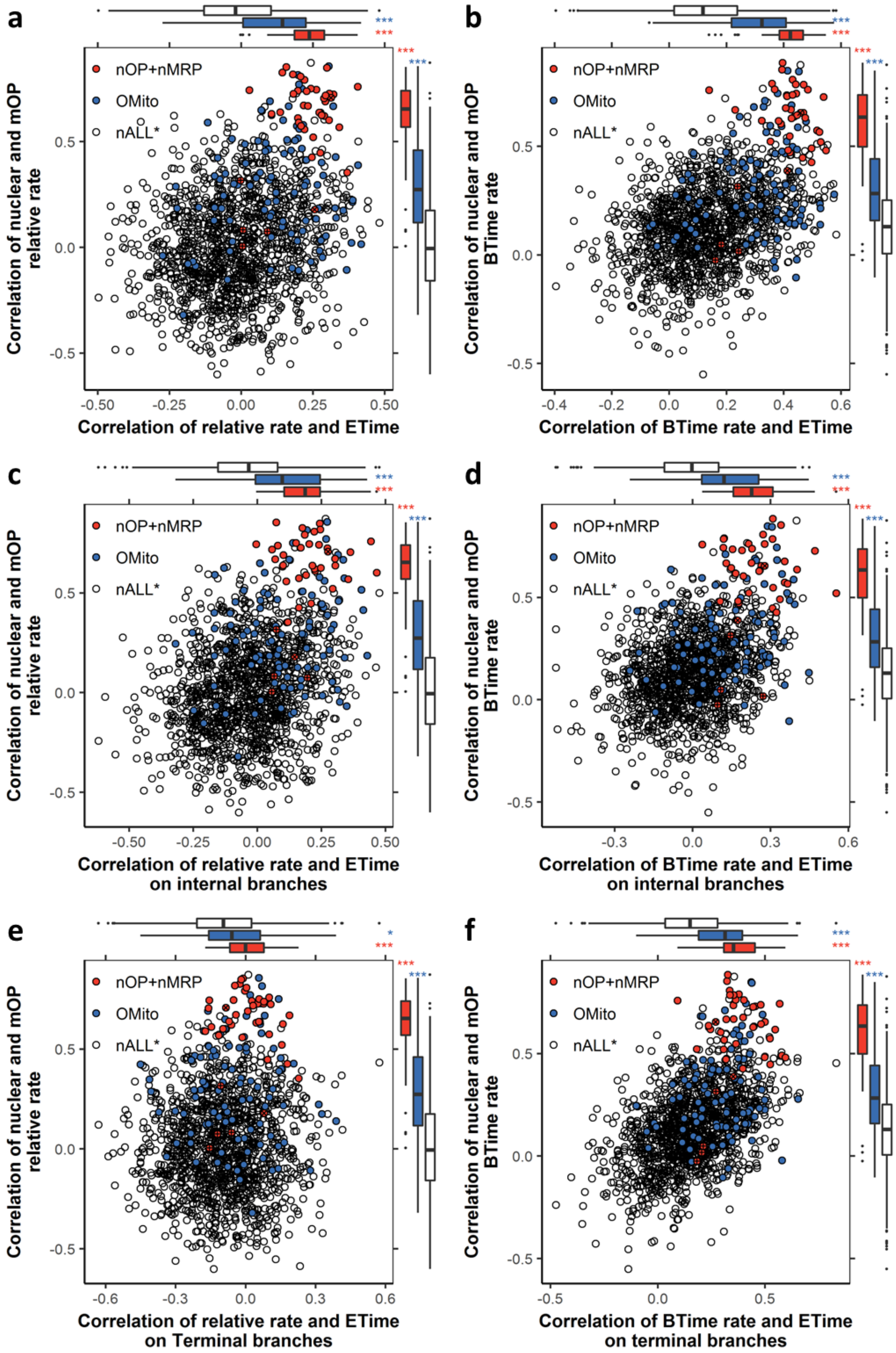


**Figure S27. Relation of evolutionary rate correlation (ERC) with mOP and apparent acceleration across evolutionary time (ETime).** Correlation of nuclear and mOP relative evolutionary rates on terminal branches and correlation of relative evolutionary rates and evolutionary time for **(a)** terminal plus internal branches, **(c)**

internal or **(e)** terminal branches only. Correlation of nuclear and mOP branch time (BTime) evolutionary rates on terminal branches and correlation of branch time rates and evolutionary time for **(b)** terminal plus internal branches, **(d)** internal or **(f)** terminal branches only. Red indicates nOP and nMRP proteins. + indicates 4 nOP proteins not in contact with mOP. × indicates 2 nOP proteins whose mOP contacting information is unknown. Blue indicate other mitochondria associated proteins identified by GO/MitoDrome annotations (OMito). Asterisks in marginal boxplot indicate significant difference from nALL*; ns indicates no significant difference ($p > 0.05$); * $p < 0.05$; ** $p < 0.01$; *** $p < 0.001$.

### Supplementary text

#### S1. Further analyses on evolutionary patterns of mitochondrial products and mitochondria-associated nuclear-encoded proteins in Holometabola

Here we provide further details on the evolutionary dynamics of both mitochondrial products (proteins and ribosomal RNAs) and mitochondria-associated nuclear-encoded proteins in Holometabola. Across holometabolous orders, we showed evolutionary rate variation for mitochondrial products (proteins and rRNA) in 44 species for which corresponding mitochondrial genomes are available (see methods). For mitochondria-associated nuclear proteins, all 64 holometabolous species in Misof *et al.* (2014) were included for stronger statistical power and a more comprehensive view of the evolution.

We initially investigated relative evolutionary rates (normalized to nALL*) on terminal branches. Variation was observed among orders for relative rates of both mitochondrial products and mitochondria-associated nuclear proteins (Figure S14; Kruskal–Wallis rank sum test (KRST); mOP: $p = 0.00045$; mrRNA: $p = 0.0035$; nOP: $p = 4.0e^{-6}$; nMRP: $p = 5.1e^{-6}$). Previous studies have reported that Hymenoptera show elevated evolutionary rates for mitochondrial genomes and nuclear-encoded OXPHOS proteins (Kaltenpoth *et al.,* 2012; Li *et al.,* 2017; Oliveira *et al.,* 2008). Consistent with this, terminal relative rates of mOP, mrRNA, nOP and nMRP all rank 1$^{st}$ in Hymenoptera among 11 investigated orders (Figure S14). In contrast, terminal relative rates of the control categories nCC and nCRP both rank 4$^{th}$ in Hymenoptera.

Figure S15 shows relative evolutionary rates for nOP, nMRP, nCC and nCRP along individual branches (both terminal and internal branches). The backbone tree topology and estimated evolutionary time were retrieved from Misof *et al.* (2014). Color indicates high (red) to low (green) relative rates along each branch. As seen in Figure S15, relative rates of both nOP and nMRP are higher on both terminal and internal branches in the Hymenoptera clade relative to other taxa (Figure S15; Wilcoxon rank sum test (WRST); nOP: $p = 4.5e^{-12}$; nMRP: $p = 1.7e^{-10}$). It appears that the elevation of evolutionary rate of nOP and nMRP in hymenopterans occurred around the time point of divergence and diversification of Hymenoptera, which is ~240 Ma (Figure S15).

The increased relative rates in Hymenoptera may be mainly caused by slow evolving of nALL*. Distance to the HCA and branch time rates on terminal branches were therefore also investigated, using the estimated divergence times for different branches in the holometabolous phylogeny (Misof *et al.*, 2014). For both mitochondrial products and mitochondria-associated nuclear proteins, their distances to HCA and terminal branch time rates vary among holometabolous orders (Figure 1, S16; KRST; $p < 0.05$ for each case). Consistent with the observation that mitochondrial evolutionary rates are elevated in Hymenoptera, mOP, mrRNA, nOP and nMRP rank 1st or 2rd in Hymenoptera for both distances to HCA and terminal branch time rates (Figure 1, S16). In contrast, the control categories nALL* and nCRP in Hymenoptera only rank from 6th to 10th.

As Hymenoptera and Diptera are both well characterized orders in Holometabola, pairwise comparisons were performed between them. For all three measurements above, Hymenoptera show a tendency for faster evolutionary rates than Diptera for mOP, mrRNA, nOP and nMRP. In contrast, Hymenoptera show slower evolutionary rates than Diptera for control nuclear categories (Figure 1, S14, S16; statistical details see below). For terminal branch relative rates, mOP, mrRNA, nOP and nMRP are significantly higher in Hymenoptera than in Diptera (Figure S14; WRST, mOP: $p$ = 8.2e$^{-5}$; mrRNA: $p$ = 8.2e$^{-5}$; nOP: $p$ = 8.0e$^{-7}$; nMRP: $p$ = 8.0e$^{-7}$), while control nuclear categories are lower but not significantly so in Hymenoptera (Figure S14). For distances to HCA, mOP, mrRNA and nOP are significantly higher in Hymenoptera than Diptera (Figure 1; WRST; mOP: $p$ = 0.0079; mrRNA: $p$ = 0.0025; nOP: $p$ = 7.8e$^{-5}$), nMRP is higher but not significantly so (WRST; $p$ = 0.13), while nuclear control categories are significant lower in Hymenoptera (Figure 1). Similarly, for terminal branch time rates, mOP, mrRNA and nOP are significant higher in Hymenoptera than Diptera (Figure S16; WRST; mOP: $p$ = 0.0016; mrRNA: $p$ = 0.0037; nOP: $p$ = 0.047), and nMRP is higher but not significantly so (WRST; $p$= 0.055), while nuclear control categories are significantly lower in Hymenoptera (Figure S16).

Therefore, we conclude that mitochondrial-nuclear coevolution rates are elevated in Hymenoptera, and in particular is higher than in Diptera. This is consistent with some observations of the role of nuclear-mitochondrial incompatibilities in species hybrids. For example, nuclear-cytoplasmic incompatibility is a major contributor to hybrid inviability in the *Nasonia* complex (Hymenoptera), but not in hybrid breakdown in *Drosophila* (Diptera) (Breeuwer & Werren 1995; Meiklejohn *et al.,* 2013; Montooth *et al.,* 2010; Niehuis *et al.,* 2008).

As already shown in the main text, nuclear-encoded proteins or amino acid sites that contact with mitochondrial-encoded products have stronger ERC with mitochondrial-encoded products than those that are not in contact. We further investigated dynamics across holometabolous orders for evolutionary rate ratios of nOP proteins in contact (**nOPcon**) over those not in contact (**nOPnot**) with mOP, and for nMRP amino acids in contact (**nMRPcon**) over those not in contact (**nMRPnot**) with mitochondrial rRNA/tRNA. Variation is observed among orders for both evolutionary rate ratio nOPcon/nOPnot and nMRPcon/nMRPnot on terminal branches (Figure S17; KRST; nOPcon/nOPnot: $p$ = 2.2e$^{-6}$; nMRPcon/nMRPnot: $p$ = 0.004). Evolutionary rate ratio nOPcon/nOPnot and nMRPcon/nMRPnot are both significantly higher in Hymenoptera than in Diptera on terminal branches (Figure S17; WRST; nOPcon/nOPnot: $p$ = 8.0e$^{-7}$; nMRPcon/nMRPnot: $p$ = 0.0085). This result suggests that higher mitochondrial evolutionary rate leads to higher evolutionary rate ratio nOPcon/nOPnot and nMRPcon/nMRPnot, which is consistent with the observation in the main text that evolutionary rate ratio for nOPcon/nOPnot and nMRPcon/nMRPnot show positive ERC with mitochondrial evolutionary rates. Thus, the result supports the conclusion that nuclear-encoded proteins or amino acid sites that contact with mitochondrial-encoded products have stronger ERC with mitochondrial-encoded products than those that are not in contact.

**S2. Mitochondrial-nuclear evolutionary rate coevolution in Psocodea**

In addition to mitochondrial-nuclear rate coevolution in Holometabola (present in main text and supplementary section S1), we also observed another independent evidence for mitochondrial-nuclear rate coevolution, within the Psocodea.

Psocodea is an order of insects, which was formerly considered as superorder. It includes what was formerly two orders, Psocoptera (barklice and booklice) and Phthiraptera (parasitic lice) (Cameron 2014). Contrasting mitochondrial substitution and gene order rearrangement rates have been reported between major lineages of Psocodea (Cameron 2014; Li *et al.,* 2015). Greatly elevated rates of mitochondrial gene order rearrangement and substitution have been observed in mitochondrial genomes from parasitic lice and booklice, but not barklice (Li *et al.,* 2013; Yoshizawa *et al.,* 2018). Mitochondrial genome fragmentation was also commonly found in parasitic lice and booklice, where a single circular mitochondrial genome separates into multiple mini-circles (Chen *et al.,* 2014).

Consistent with the idea that evolutionary rates co-evolve between mitochondrial products and mitochondria-associated nuclear proteins, we also observed elevated evolutionary rate of nOP and nMRP within Psocodea. We did not investigate mitochondrial evolutionary rates within Psocodea, as we were only able to retrieve mitochondrial genomes (in the same families) corresponding to 2 of 4 Psocodea species in the Misof *et al.* (2014) tree.

Among 32 investigated orders in Hexapoda, Psocodea ranks 1$^{st}$ for terminal relative rates of both nOP and nMRP (Figure S18a–b). In addition, acceleration of mitochondria-associated nuclear-encoded proteins in Psocodea seems to occur after the divergence from *Ectopsocus briggsi* (barklice, Psocoptera, Psocomorpha), which is estimated ~187 Ma (Figure S18c–d). As there are only 4 Psocodea species and not enough for effective statistical comparison for concatenated sequences, we compared their evolutionary rates on terminal branches at the individual gene level (Figure S19). Compared to *E. briggsi* (barklice), proteins in nOP and nMRP both show significantly higher terminal relative rates in *Liposcelis bostrychophila* (booklice, Psocoptera, Troctomorpha), *Menopon gallinae* (parasitic lice, Phthiraptera) and *Pediculus humanus* (parasitic lice, Phthiraptera) (Figure S19a; Wilcoxon signed-rank test; $p <$ 0.001 for each case). In contrast, proteins in nuclear control categories nCC and nCRP don't show these patterns, except the terminal relative rate of nCRP is significantly higher in *L. bostrychophila* than in *E. briggsi* (Wilcoxon signed-rank test; $p < 0.05$).

GO enrichment analysis was further performed for the top 10 % of proteins with fastest relative rates in each Psocodea species. Several mitochondria-related "Cellular Component" items were overrepresented in the top 10% fastest relative rates proteins in *L. bostrychophila*, *M. gallinae* and *P. humanus*, but not in *E. briggsi* (Figure S19b; 13 mitochondria-related of 25 items for *L. bostrychophila*, 12 of 32 for *M. gallinae*, 12 of 27 for *P. humanus*, 0 of 22 for *E. briggsi*; Fisher's exact test between *E. briggsi* and other three species, $p = 1.3e^{-5}$). No mitochondria-related "Cellular Component" items were overrepresented in the bottom 10% of slowest relative rate proteins in any of the four species (data not shown).

These results provide evidence of another order (Psocodea) in which mitochondrial evolutionary rates are elevated, with a corresponding elevation in mitochondrial associated nuclear-encoded proteins.

#### S3. Apparent acceleration of nuclear proteins associated with mitochondria

Apparent acceleration has also been reported in metazoan mitochondrial genes, where substitution rate estimates are higher in recent timescales than for those in older timescales (Ho *et al.*, 2005; Molak & Ho 2015). However, incomplete purifying selection at shallow branches and saturation at deep branches were proposed by authors to explain this pattern (Molak & Ho 2015). Here we present analyses of changes in evolutionary rates of mitochondrial-associated proteins over evolutionary time, to determine if they also show apparent acceleration in evolutionary time. The basic question is "Are mitochondrial proteins and mitochondria-associated nuclear-encoded proteins accelerating their rates of change over evolution?".

##### S3.1 Correlation between branch time rates and evolutionary time

Based on the relative evolutionary rates mapped to Misof *et al.* tree (Figure S15), we observed an apparent acceleration for mitochondria-associated nuclear-encoded proteins across evolutionary time in holometabolous insects. To test this observation, correlation analyses were performed between evolutionary rates of branches and their evolutionary time, which is the time span from tip to midpoint of branches.

We first focus on changes of branch time rates across evolutionary time, because this allows a comparison of time-dependent patterns in mitochondria-associated versus other nuclear proteins. Non-parametric Spearman correlations were conducted between branch time rates and calibrated evolutionary time in Holometabola. Branch time rates of nALL*, **nOPnMRP** (combined nOP and nMRP), nOP, nMRP and nCRP show significantly positive correlation with evolutionary time, while nCC doesn't (Figure 5, S20; nALL*: $\rho = 0.29$, $p = 0.001$; nOPnMRP: $\rho = 0.60$, $p = 8.3e^{-14}$; nOP: $\rho = 0.55$, $p = 1.6e^{-11}$; nMRP: $\rho = 0.60$, $p = 1.4e^{-13}$; nCRP: $\rho = 0.27$, $p = 0.002$;

nCC: $p = 0.08$). Although all tested nuclear categories, except nCC, show apparent acceleration, branch time rate of nOPnMRP, nOP and nMRP show significantly higher correlations with evolutionary time than those of nuclear control categories nALL*, nCRP and nCC (dependent Spearman correlations comparison; $p < 0.001$ for each case).

As incomplete purifying selection was proposed to explain time-dependent evolutionary rate pattern observed in mitochondrial genomes (Ho *et al.*, 2005; Molak & Ho 2015), apparent acceleration patterns of mitochondria-associated nuclear-encoded (and other) proteins may be mainly caused by incomplete purifying selection. We next examined apparent acceleration of branch time rates on terminal branches. On terminal branches, branch time rates of nALL*, nOPnMRP, nOP, nMRP, nCRP and nCC all show significantly positive correlations with evolutionary time (Figure 5, S20; nALL*: $\rho = 0.44$, $p = 0.0003$; nOPnMRP: $\rho = 0.55$, $p = 2.2e^{-6}$; nOP: $\rho = 0.51$, $p = 2.0e^{-5}$; nMRP: $\rho = 0.57$, $p = 9.5e^{-7}$; nCRP: $\rho = 0.28$, $p = 0.026$; nCC: $\rho = 0.32$, $p = 0.009$). In other words, longer terminal branches have lower apparent evolutionary rates. Branch time rates were also compared between terminal branches and their matched parent internal branches, confirming terminal branches show higher branch time rates than their parent internal branches for nALL*, nOPnMRP, nOP and nMRP, but not for nCRP and nCC (Wilcoxon signed rank test; nALL*: $p = 0.005$; nOPnMRP: $p = 0.0008$; nOP: $p = 0.02$; nMRP: $p = 0.0002$; nCRP: $p = 0.055$; nCC: $p = 0.24$). These results suggest standing variation at “tips” (i.e. incomplete purifying selection) may contribute to terminal branch rate estimation and create an apparent time-dependent evolutionary rate pattern.

To reduce the incomplete purifying selection effect, we investigated time dependent rate pattern only on internal branches. The reasoning is that contributions of standing variation at the terminal branches would be absent in the internal branches. The minimum and median evolutionary time span of terminal branches in Holometabola are 27.34 and 101.27 Ma, respectively. These deep terminal branches left internal branches unlikely to be influenced by incomplete purifying selection. For internal branches, only branch time rates of nOPnMRP, nOP and nMRP still show significantly positive correlations with evolutionary time (Figure 5, S20; nOPnMRP: $\rho = 0.41$, $p = 0.001$; nOP: $\rho = 0.45$, $p = 0.0003$; nMRP: $\rho = 0.39$, $p = 0.002$), while nALL*, nCC and nCRP do not (Figure 5, S20; nALL*: $p = 0.96$; nCRP: $p = 0.99$; nCC: $p = 0.68$). This suggests that apparent acceleration of nuclear control categories (i.e. nALL* and nCRP) may be mainly explained by incomplete purifying selection on terminal “tips”, but apparent acceleration of mitochondria-associated nuclear proteins (i.e. nOPnMRP, nOP and nMRP) cannot be easily explained by incomplete purifying selection.

To reduce the contribution of saturation as a potential cause, subsampling analyses were conducted. After controlling evolutionary rates of proteins or conservation scores of sites, branch time rates of nOPnMRP, nOP and nMRP all show significantly higher positive correlation with evolutionary time than randomly generated samples (Figure S11, S21a–b, $p < 0.001$ for each case). Results were similar for branch time rates on internal or terminal branches only (Figure S21c–f). These indicate that saturation is unlikely to be the cause of apparent acceleration for mitochondria-associated nuclear proteins. Some other features of nOPnMRP, nOP and nMRP (gene number size, alignment length and gap percentage) are also tested, and unlikely to be the causes either.

In addition, results were confirmed after removing Hymenoptera (Figure S22), ruling out that apparent acceleration patterns are mainly caused by Hymenoptera evolutionary acceleration in mitochondria-associated nuclear proteins.

Because recessive lethal and deleterious genes are exposed to selection in haploid males, it has been widely recognized that the equilibrium frequencies of such alleles will be lower in haplodiploid than diploid species (Werren 1993), and hence incomplete purifying selection on terminal branches less of a factor. We therefore investigated evolutionary time-dependent branch rate patterns using data from Peter *et al.* (2017) on hymenopteran phylogeny and protein alignments. Strikingly, branch time rates of nALL* and nCC show a significantly negative correlation with evolutionary time (Figure 5, S23; nALL*: $\rho = -0.22$, $p = 4.7e^{-5}$; nCC: $\rho = -0.24$, $p = 8.9e^{-6}$), and branch time rate of nCRP shows no significant correlation with evolutionary time ($p = 0.46$). In contrast, branch time rates of nOPnMRP, nOP and nMRP show significantly positive correlations with evolutionary time in Hymenoptera (Figure 5, S23; nOPnMRP: $\rho = 0.38$, $p = 5.3e^{-13}$; nOP: $\rho = 0.32$, $p = 2.3e^{-9}$; nMRP: $\rho = 0.37$, $p = 4.9e^{-12}$). Similar patterns were also confirmed when using only terminal or internal branches (Figure S23). The results for nALL* are consistent with the idea of reduced standing variation in Hymenoptera on terminal branches (e.g. due to more effective purifying selection) (Figure 5). The patterns are now quite distinct between nALL* and the mitochondria-associated nuclear-encoded proteins (nOP and nMRP), which suggest that they have very different histories over evolutionary time.

However, for all described correlations between branch time rates and evolutionary time in this section, no significant correlations held after phylogenetic correction using Bayesian mixed model (Table S5–6, 8) (Hadfield & Nakagawa 2010). Therefore, we conclude that mitochondria-associated nuclear proteins show strikingly different time-dependent branch time rate patterns from other nuclear-encoded proteins. But there is no clear evidence supporting an acceleration of mitochondria-associated nuclear proteins over evolutionary time.

##### S3.2 Correlation between relative evolutionary rates and evolutionary time

In addition to time-dependent analyses of branch time rate, we also investigated changes of relative evolutionary rate (normalized to nALL*) across evolutionary time.

Relative rates of nOPnMRP, nOP and nMRP show significantly positive correlations with evolutionary time (Figure 5, S24; nOPnMRP: $\rho = 0.30$, $p = 0.0006$; nOP: $\rho = 0.29$, $p = 0.001$; nMRP: $\rho = 0.30$, $p = 0.0007$). In contrast, relative rate of nCC shows significantly negative ($\rho = -0.23$, $p = 0.0089$), and nCRP shows no significant correlation with evolutionary time ($p = 0.26$). To reduce the incomplete purifying selection effect, we also investigated time-dependent rate pattern only on internal branches. For internal branches, relative evolutionary rates of nOPnMRP, nOP and nMRP still show significantly positive correlation with evolutionary time, but control categories show no significant correlation (Figure 5, S24; nOPnMRP: $\rho = 0.41$, $p = 0.001$; nOP: $\rho = 0.45$, $p = 0.0003$; nMRP: $\rho = 0.39$, $p = 0.002$; nALL*: $p = 0.96$; nCRP: $p = 0.99$; nCC: $p = 0.68$). Correlations and significances held after phylogenetic correction using Bayesian mixed model (Table S5). However, it should be noted that changes in relative rate over time can be due to changes in the numerator (e.g. nOP branch time rate) or denominator (i.e. nALL* branch time rate) or a combination of both.

To reduce the contribution of saturation as a potential cause, subsampling analyses were conducted. After controlling evolutionary rates of proteins or conservation scores of sites, relative rates of nOPnMRP, nOP and nMRP show significantly higher positive correlation with evolutionary time than randomly generated samples (Figure S11, S25a–b; $p < 0.05$ for each case). Similar patterns and significances were confirmed on internal branches only but not on terminal branches (Figure S25). These results indicate that saturation cannot easily explain apparent acceleration of relative rates in nOPnMRP, nOP or nMRP. Some other features of nOPnMRP, nOP and nMRP (gene number size, alignment length and gap percentage) are also tested, and unlikely to be the causes either.

In addition, similar results were found after removing Hymenoptera (Figure S22, Table S8), excluding Hymenoptera as the cause of the apparent evolutionary acceleration.

We also examined the expanded set of Hymenoptera using data from Peters *et al.* (2017) for patterns in relative evolutionary rate compared to evolutionary time midpoint for each branch. Relative rates of nOPnMRP, nOP, nMRP and nCRP all show significantly positive correlation with evolutionary time (Figure 5, S26; nOPnMRP: $\rho = 0.47$, $p = 4.5e^{-20}$; nOP: $\rho = 0.43$, $p = 2.8e^{-16}$; nMRP: $\rho = 0.47$, $p = 1.1e^{-19}$; nCRP: $\rho = 0.20$, $p = 0.0002$). In contrast, relative rate of nCC shows significantly negative correlation with evolutionary time ($\rho = -0.20$, $p = 0.0002$). Correlations and significances held after phylogenetic correction. For internal branches, relative evolving rates of nOPnMRP, nOP and nMRP still show significantly positive correlation with evolutionary time, but nCC shows significantly negative correlation and nCRP shows no significant correlation with evolutionary time (Figure 5, S26; nOPnMRP: $\rho$ = 0.49, $p$ = 1.2e$^{-11}$; nOP: $\rho$ = 0.44, $p$ = 3.9e$^{-9}$; nMRP: $\rho$ = 0.50, $p$ = 4.7e$^{-13}$; nCRP: $p$ = 0.065; nCC: $\rho$ = -0.18, $p$ = 0.020). However, for relative rates on internal branches, correlations and significances didn't hold after phylogenetic correction (Table S6).

These results suggest that relative rates of mitochondria-associated nuclear-encoded proteins increase across evolutionary time (i.e. apparent acceleration). However, it may be caused by artifacts, for example branch length estimation or rate normalization. Furthermore, using relative rates (i.e. normalization to nALL* to determine rates) obscures the role of changes in nALL* protein rates over evolutionary time.

##### S3.3 Individual gene analysis for apparent acceleration of mitochondria-associated nuclear-encoded proteins

To have more comprehensive understandings of time-dependent evolutionary rate patterns, individual gene analyses were conducted. To treat every gene equally, the concatenated set of 1478 individual genes was used for relative rate normalization. nALL* was also tested for relative rates normalization, and showed with similar patterns to those described below.
Figure S27 plots the correlation of nuclear protein evolutionary rates to evolutionary time, versus evolutionary rate correlations (ERC) between nuclear proteins and mOP. Highly significantly positive correlations are found (Figure S27a, relative rate: $\rho$ = 0.26, $p$ = 4.3e$^{-25}$; Figure S27b, branch time rate: $\rho$ = 0.38, $p$ = 5.4e$^{-51}$). This suggests a potential unknown relationship between mitochondria-association and apparent acceleration of evolutionary rate. In addition, both branch time rates and relative rates of nOPnMRP show higher correlations with evolutionary time (i.e. apparent acceleration) than do nALL* ($p < 0.001$), which is consistent with results using concatenated proteins. We also observed that both branch time rate and relative rate of OMito show higher ERC with evolutionary time (i.e. apparent acceleration) than do nALL* ($p < 0.001$). Similar results were also confirmed for both relative and branch time rate only on terminal or internal branches. In each case, nOPnMRP and OMito both show significantly higher correlations with evolutionary time than do nALL* (Figure S27, $p < 0.05$ for each case). Therefore, individual gene analysis also indicate different time-dependent evolutionary rate patterns for mitochondria-associated versus other nuclear-encoded proteins. Specifically, evolutionary rate of mitochondria-associated nuclear proteins are more likely to show apparent acceleration than other nuclear-encoded proteins.

#### Supplementary methods

##### Phylogenetic analysis in Hymenoptera using Peters *et al.* (2017) data

To reduce the effect of incomplete purifying selection on apparent acceleration, time-dependent evolutionary patterns were investigated in Hymenoptera using Peters *et al.* (2017) phylogeny and alignments. Similar to the analysis in Holometabola using Misof *et al.* data, protein alignments of the 3256 single copy nuclear genes and the dated tree were retrieved from supplementary materials of Peters *et al.* (2017).

For defining nuclear-encoded protein categories, GO and KEGG annotations were determined based on their *Apis mellifera* orthologs. Among 3256 single copy genes from *A. mellifera*, 2884 have *Drosophila* orthologs in OrthoDB v7 (Zdobnov *et al.*, 2017), and their GO annotations were retrieved from FlyBase v2017_06 (Gramates *et al.*, 2017). For the remaining 372 *A. mellifera* proteins without *Drosophila* orthologs, GO annotations were assigned based on their motif and domain information using InterProScan v5.2 (Jones *et al.*, 2014). For KEGG annotation, *A. mellifera* proteins were annotated using the BlastKOALA online service (Kanehisa *et al.*, 2016).

Based on GO or KEGG information, nuclear protein categories were defined using the same method described for Misof *et al.* data. Briefly, 248 proteins were identified as mitochondria-associated proteins, using GO or MitoDrome information. Among these 248 proteins, 27 were identified as nuclear-encoded oxidative phosphorylation proteins (nOP); 39 were identified as nuclear-encoded mitochondrial ribosomal proteins (nMRP). The remaining 3008 nuclear proteins (all 3256 single copy proteins minus 248 mitochondria-associated proteins) were defined as nALL*. Among these nALL* proteins, 32 proteins were identified as nuclear-encoded cell cycle proteins (nCC); 34 were identified as nuclear-encoded cytosolic ribosomal proteins (nCRP). For these nuclear protein categories, original alignments were first filtered by trimAI version 1.2rev59 with automated settings (Capella-Gutierrez *et al.*, 2009), and then concatenated using AMAS (Borowiec 2016). Branch lengths were estimated based on the Peters *et al.* (2017) tree using RAxML v8.0.20 by setting substitution model as "PROTGAMMAIAUTO" (Stamatakis 2014). Branch lengths and corresponding branch times were extracted using Newick Utilities v1.6 (Junier & Zdobnov 2010).
